# Elevated hydrostatic pressure acts via piezo-1 to destabilise VE-cadherin junctions: an endothelium-on-chip study

**DOI:** 10.1101/2025.03.27.645710

**Authors:** P. Vasanthi Bathrinarayanan, Thomas Abadie, Patricia Perez Esteban, D. Vigolo, M. J. H. Simmons, L. M. Grover

## Abstract

Despite the effects of shear stress on endothelial biology having been extensively researched, the effects of hydrostatic vascular pressure at extremely low shear stresses have been largely ignored. In the current study, we employ a microfluidic organ-on-chip platform to elucidate the time and shear stress dependent effects of elevated hydrostatic pressure on endothelial junctional perturbations. We report that short term (1h) exposure to elevated hydrostatic pressure at high shear stress (1 dyne/cm^2^) but not low shear stress (0.1 dyne/cm^2^) caused VE-cadherin to form serrated, finger like projections at the cell-cell junctions and this effect was abrogated upon pharmacologically inhibiting piezo-1 mechanosensory protein. Interestingly, prolonged exposure (24h) to elevated hydrostatic pressure at low (0.1 dyne/cm^2^) but not high shear stress (1 dyne/cm^2^) caused VE-cadherin internalisation, thereby increasing the cytoplasmic concentration. Further, we report that this internalisation of VE-cadherin was reversible upon pharmacologically inhibiting piezo-1 in a time-dependent manner wherein after 12h, we observed stable VE-cadherin junctions re-appear at the cell-cell junctions. Overall, we demonstrate that piezo-1 plays a crucial role in the mechanotransduction of elevated hydrostatic pressure by regulating the VE-cadherin dynamics at cell-cell junctions. Targeting piezo-1 may provide a novel therapeutic/diagnostic marker, especially in conditions that involve microvascular dysfunction due to elevated vascular pressures.

## 1 Introduction

Cells dynamically respond to biochemical factors and a diverse range of mechanical forces that influence critical cellular processes such as differentiation, proliferation, stiffness, and migration ^1, 2^. The vasculature, in particular, experiences three distinct mechanical forces- shear stress, circumferential stretch and intraluminal hydrostatic pressure. The role of shear stress in facilitating endothelial dysfunction has been well established ^3^. In comparison, although abnormal elevations in hydrostatic pressure are a major contributor to several diseases like glaucoma ^4^, traumatic brain injury (TBI) ^5^ and pulmonary oedema ^6^, very sparse information is available on the mechanobiological role of hydrostatic pressure in vascular disease progression. Acute compartment syndrome (ACS) is one such condition wherein a sudden increase in the intracompartmental pressure due to haemorrhage or oedema can cause rapid elevations in microvascular pressure, restricting blood flow into the tissue ^7^. If left untreated, the collapsed microvasculature can cause ischemia ^8^ and microvascular dysfunction which results in permanent neuro-muscular damage ^9^. The mechanisms by which endothelial cells sense and respond to increases in hydrostatic pressure are largely unknown, and hence, investigations into these phenomena could lead to novel diagnostic and therapeutic interventions for the benefit of ACS patients.

Junctional contacts in endothelial cells are predominantly mediated by vascular endothelial cadherins (VE-cadherin) in conjunction with other adhesion proteins such as occludins, claudins, nectins, JAMs and PECAM-1 ^10^. VE-cadherin (also known as CD144) is a transmembrane adherens junction protein that has been extensively studied for its role in regulating vascular junctional integrity ^11–13^. Past studies have shown that changes in shear stress can significantly alter VE-cadherin dynamics and, hence, influence permeability, coagulation and inflammation ^14, 15^. In comparison, very few studies ^16–18^ have reported on the influence of isotropically applied hydrostatic pressure on VE-cadherin junctional dynamics, especially in response to the contractile forces generated by the actomyosin cytoskeleton which can lead to VE-cadherin disruption at cell-cell junctions ^19^. These past studies, although very informative, consisted of static cultures and did not study the combined influence of hydrostatic pressure at different flow induced shear stresses. Importantly, since low shear stresses (<5 dyne/cm^2, 20^) have been implicated in the pathophysiology of many vascular conditions ^21, 22^, the influence of elevated hydrostatic pressure at low shear stresses would shed light on previously unknown endothelial mechanotransduction mechanisms. Toward this goal, this study investigated the combined influence of shear stress and elevated hydrostatic pressure on VE-cadherin and actin dynamics using a microfluidic model of a blood vessel.

## 2 Experimental

### 2.1 Cell culture

Primary human umbilical vein endothelial cells (HUVECs) were chosen for this study as they faithfully represent the in vivo human endothelium behaviour compared to the other endothelial cell lines ^23^. Owing to its physiological relevance, HUVECs have been extensively used in both academia as well as biopharmaceutical industries to investigate vascular pathologies ^24^. HUVECs (PromoCell, Heidelberg, Germany) between passages 3-8 were cultured in M199 growth medium supplemented with 10% fetal bovine serum (FBS) (Gibco™, Sigma-Aldrich, UK) and 15mL endothelial cell growth supplement mix (Cell applications Inc., Merck, UK) per 500mL media bottle.

### 2.2 Microfluidic experiments

Ibidi µ-Slide VI 0.1 (Ibidi GmbH, Gräfelfing, Germany) microfluidic channels (1 mm wide, 100 µm height and 17 mm length) were used for the microfluidic experiments. Prior to seeding, the channels were coated with 0.2% gelatine solution (Sigma Aldrich, Merck Ltd, Dorset, UK) and incubated at 37°C and 5% CO_2_ for 2h. Channels were washed with fresh growth medium and HUVECs were gradually introduced into the channels at a density of 10^5^ cells/channel (i.e. 10 µL of HUVECs from a cell suspension of 10^7^ cells/mL) without forming air bubbles. Cells were observed under the brightfield microscope (EVOS M3000, Invitrogen, Fisher Scientific, Loughborough, UK) to ensure that there was no movement of the cells as this would prevent their adhesion to the channel surface. 60 µL of growth media was added to the inlet and the outlet ports of the channel after which the cells were incubated at 37°C and 5% CO_2_ for 2 h. After 2 h, HUVECs formed a uniform monolayer on the channels and were now ready for perfusion.

### 2.3 Shear stress and pressure distributions

Syringe pumps (World Precision Instruments, Hitchin, UK) were used to achieve the desired shear stress (SS) for different conditions by adjusting the flow rate (*Q*). For a rectangular channel with width 2𝑊 and height 2*H*, the velocity field can be written ^25^:

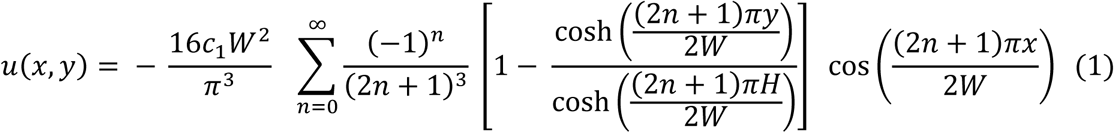

Where *c_1_* depends on the average velocity, 𝑢_𝑚_ = 𝑄/(2𝑊 × 2𝐻), and can be written as:

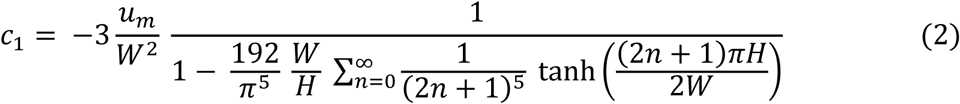

From this, the wall shear stress (WSS) at the bottom wall of the channel (where the cells are cultured) is written as:

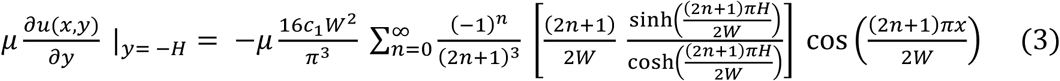

Figure 1 (A) and (B) represent the schematic of the microfluidic channel set-up during normal and elevated pressure flow conditions respectively. The WSS distribution along the bottom wall of the channel and the corresponding velocity counter plots across the channel cross section for both flow rates, namely 1.3 μL/min for low shear stress and 13 μL/min for high shear stress experiments, are depicted in Figure 1 (C,C_i_) and Figure 1 (D,D_i_) respectively. Both conditions demonstrated a uniform shear stress profile with a peak WSS value of 0.014 Pa (0.14 dyne/cm^2^) and 0.14 Pa (1.4 dyne/cm^2^) for the low and high flow rates, respectively.

**Figure 1.**
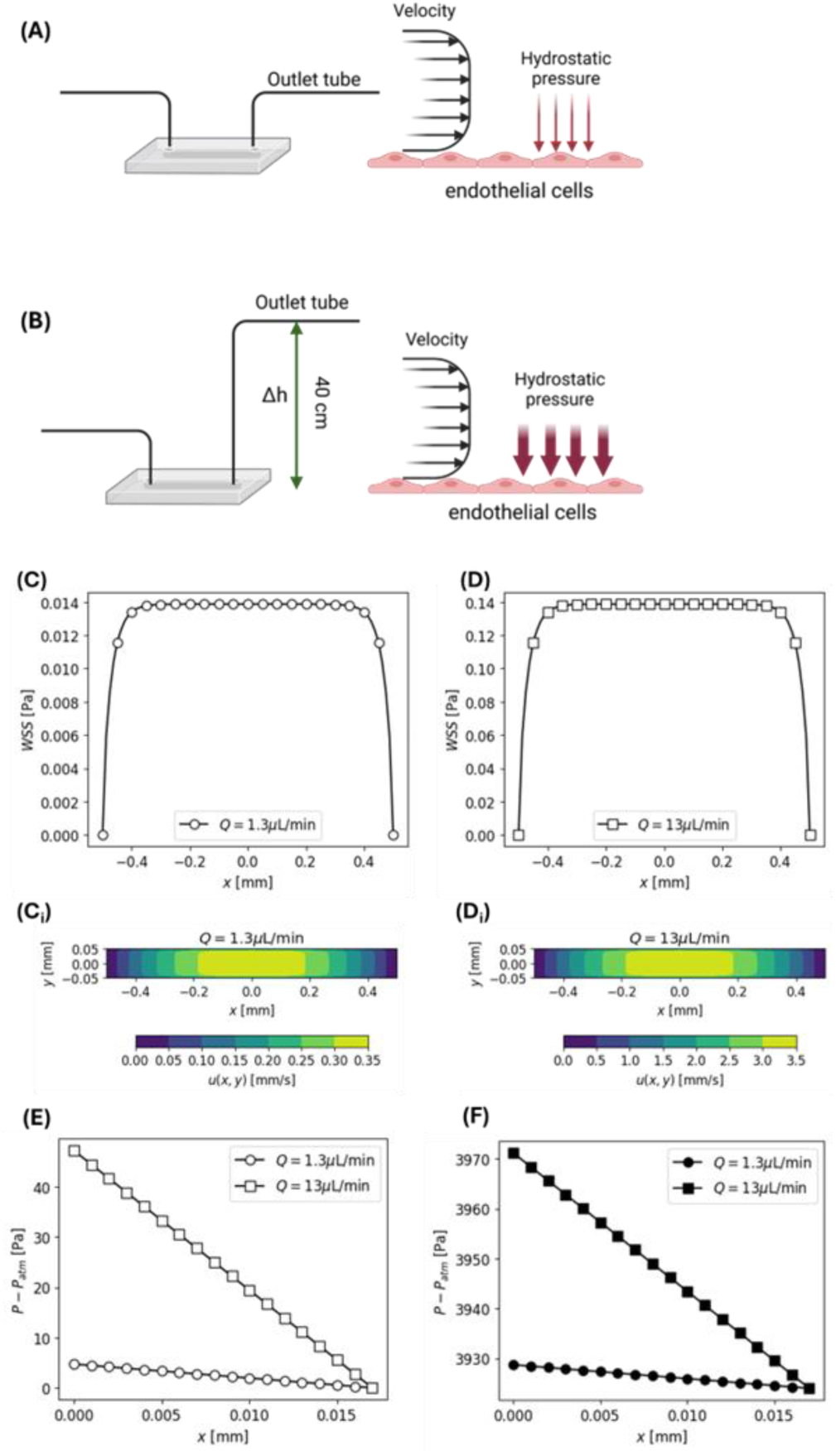
Wall shear stress (WSS) and pressure distribution within the channel. (A) Schematic of the microfluidic channel perfusion during normal flow conditions and (B) during application of elevated hydrostatic pressure conditions during which the height of the outlet tube was elevated by 40 cm relative to the inlet tube, thereby maintaining the same shear stress profile but increasing the pressure experienced by the cells. (C,D) WSS distribution at the channel bottom surface derived from the velocity counter plots (C_i_,D_i_) for the low flow rate and high flow rate respectively. (E) Pressure distribution along the channel length for the low and high flow rates under normal conditions and (F) at elevated hydrostatic pressure conditions.

Elevated hydrostatic pressure (HP) conditions were achieved by raising the outlet tube to 40 cm above the inlet as shown in Figure 1 (B). At this height, the pressure drop at the channel outlet (𝑃_𝑜𝑢𝑡_) can be derived from:

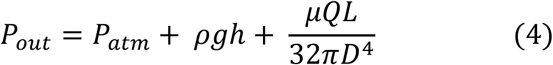

where 𝑃_𝑎𝑡𝑚_is the atmospheric pressure, 𝜌 and 𝜇 are the density (1,000 kg m^-3^) and dynamic viscosity (0.00072 Pa s at 37°C) of the growth medium, *h* is the height of the outlet tubing (40 cm), *Q* is the flow rate and *D* is the inner diameter (1 mm) of the outlet tubing. Given the very low flow rates, the friction pressure drop is negligible (<1 Pa) when compared to the atmospheric pressure. Hence the pressure drop at the outlet is 𝑃_𝑜𝑢𝑡_ − 𝑃_𝑎𝑡𝑚_ = 3924 𝑃𝑎 (29.43 mmHg). In conditions such as ACS, an intracompartmental pressure ≥ 30 mmHg is used as a clinical diagnostic criterion ^26^, and hence, we wanted to investigate the impact of this elevated pressure on the endothelial cells.

For a rectangular channel, the pressure drop along the channel can be expressed ^25, 27^:

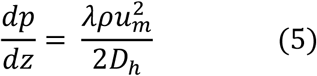

where 𝐷_ℎ_ is the hydraulic diameter and *λ* depends on the aspect ratio of the channel (𝛼 = 𝑊/𝐻) and the Reynolds number (𝑅𝑒 = 𝜌𝑢_𝑚_𝐷_ℎ_/𝜇):

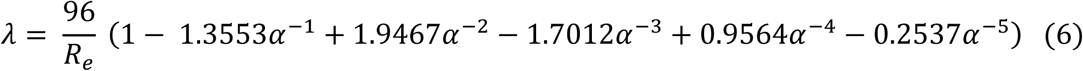

The pressure at a given point in the channel, with *P_out_* being the pressure at the exit of the channel, *L* being the channel length (L=17 mm) and *z* being the position from the inlet (i.e. *L−z* is the position in the channel relative to the outlet) is given by:

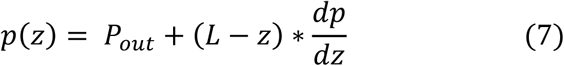

The pressure difference (ΔP) across the microfluidic channel was 4.7 Pa to 47 Pa for flow rates 1.3 μL/min and 13 μL/min under normal and elevated hydrostatic pressure conditions respectively as shown in Figure 1 (C) and (D). A summary of the two flow rates used, shear stress and pressure at the inlet, is presented in Table 1.

**Table 1.** Flow rates for different test conditions and their associated shear stress and pressure head (at inlet) values.

| | Flow rate ($\mu\text{L}/\text{min}$) | Wall shear stress (dyne/cm<sup>2</sup>) | Pressure (Pa) at inlet (~) |
| --- | --- | --- | --- |
| SS high only | 13 | 1.4 | 47.1 |
| SS low only | 1.3 | 0.14 | 4.71 |
| SS high+HP | 13 | 1.4 | 3971.1 |
| SS low+HP | 1.3 | 0.14 | 3928.7 |

### 2.4 Piezo-1 inhibition

For piezo-1 inhibition experiments, cells were exposed to growth media that contained GsMTx4 (Tocris bioscience, Bio-Techne, UK) which is a peptide derived from tarantula venom that selectively inhibits mechanosensitive cationic channels including piezo-1 ^28,29^. GsMTx4 was dissolved in growth media to give a final working solution of 5 µM concentration. After exposure, cells were washed thrice with DPBS and were fixed for immunostaining.

### 2.5 Immunofluorescence staining

For immunofluorescent staining, the cells were fixed by flowing 10% Formalin neutral buffered solution (Sigma Aldrich, Merck Ltd, Dorset, UK) for 10 min at room temperature into the microchannel followed by incubation with 0.1% Triton-X in Dulbecco’s phosphate buffered saline (DPBS, Gibco, Merck Ltd, Dorset, UK) for 5 min at room temperature for permeabilization of the cell membrane. The channels were then washed 3 times using DPBS and were subsequently blocked using 1% BSA (Sigma Aldrich, Merck Ltd, Dorset, UK) diluted in DPBS containing 0.1% Tween-20 (Sigma Aldrich, Merck Ltd, Dorset, UK). The channels were washed thrice with DPBS and were incubated with 5 µg/mL (diluted in blocking buffer) mouse anti-human monoclonal VE-cadherin (CD144) primary antibodies (Invitrogen, Fisher Scientific, Loughborough, UK) at 4°C overnight. The following day, the unbound primary antibodies were removed by washing the channels three times with DPBS and the cells were subsequently incubated for 1 h at room temperature with goat anti-mouse secondary antibodies conjugated with Alexa Fluor™ 488 (Invitrogen, Fisher Scientific, Loughborough, UK) at a dilution of 1:1000 in blocking buffer. After 1 h, the unbound secondary antibodies were removed by washing 3 times with DPBS. The cells were then stained for F-actin using Phalloidin CF®594 (Biotium Inc., CA, USA) as well as nuclei using DAPI (4’,6-diamidino- 2-phenylindole, Sigma Aldrich, Merck Ltd, Dorset, UK) for 5 min at room temperature. The cells were then imaged using a Carl Zeiss LSM880 scanning confocal microscope predominantly using the 40x (C-Apochromat, 1.2 W Korr FCS M27) and 63x (C-Apochromat, 1.2 W Korr M27) objectives.

### 2.6 Image analysis

All the raw images were processed (brightness/contrast adjustments) either using FIJI (US National Institutes of Health) or Zeiss Zen software. Cells were predominantly imaged around the middle area of the channel cross-section where peak WSS was experienced. Cells at the very edges of the channel (growing on the side walls) and cells around the inlet/outlet ports were avoided. For VE-cadherin and actin co-localisation plot profiles on FIJI, a straight line was drawn on either a remodelling junction or a continuous junction. For all other cell morphological analyses, customised pipelines on CellProfiler 4.2.6 ^30^ (Broad Institute, Massachusetts Institute of Technology, USA) were used to automate image analysis. Briefly, nuclei and actin raw images, designated as primary and secondary objects, respectively, were thresholded using a minimum cross-entropy method. Cytoplasm fluorescence intensity was derived by subtracting the primary object (DAPI stain) from the secondary object (whole cell stain, which in this case is actin stain). Cell area, circularity, compactness, and whole cell VE-cadherin fluorescence intensity were quantified using the ‘Measure ObjectSizeshape’ and ‘Measure ObjectIntensity’ modules. Membrane VE-cadherin fluorescence intensity was measured using a modified pipeline. Briefly, the primary and secondary objects were obtained using the DAPI and actin images. The secondary object was then shrunk by 3 pixels using the ‘ExpandorShrinkObjects’ module. The cell membrane of 3 pixel thickness was isolated by subtracting the shrunk secondary object from the whole secondary object and the VE-cadherin fluorescence intensity was measured using the ‘Measure ObjectIntensity’ module. The pipelines used for image analysis on CellProfiler can be provided upon request.

### 2.7 Statistics

All statistical analysis was performed using GraphPad Prism10.3.1. One-way ANOVA along with Tukey’s post-hoc test was used to compare between different conditions. Statistical significance was set as follows: *p < 0.05, **p < 0.01, ***p < 0.001****p < 0.0001. Statistical tests and relative p values are indicated in each figure legend. Unless stated elsewhere, all experiments were performed with at least three biological independent (3 channels) replicates.

## 3 Results

### 3.1 Elevated hydrostatic pressure produces VE-cadherin finger like projections at junctions

To test the short-term influence of elevated hydrostatic pressure at low and high shear stresses, cells within the microfluidic channels were exposed to 0.14 dyne/cm^2^ or 1.4 dyne/cm^2^ for 1h after which cells were fixed and stained for VE-cadherin, actin and nuclei. Cells in all three conditions demonstrated a polygonal morphology with no difference in eccentricity although the cells exposed to SS high+HP conditions demonstrated a more irregular shape compared to the other conditions as shown in Figure 2 (A-D). All three conditions exhibited thick, parallelly aligned stress fibres that traversed the entire cell body compared to the cortical actin fibres that appeared in control cells. There was no particular orientation of actin fibres in the direction of flow with different areas of the channels demonstrating different orientation angles (refer Supplementary Figure S1 for orientation angles at different locations of the channel). This lack of alignment is possibly due to the short exposure time of 1 h as endothelial cells generally take between 6-24 h to exhibit alignment and elongation in the direction of the flow ^31, 32^. Cells exposed to SS high+HP demonstrated highest relative VE-cadherin fluorescence intensity (F.I.) in both whole cells as well as at the cell membrane as depicted in Figure 2 (c,d). In contrast, SS low+HP demonstrated the lowest relative VE-cadherin F.I., suggesting that an application of hydrostatic pressure at high shear stress causes an increase in junctional VE-cadherin presence, although the pattern of expression can vary.

**Figure 2.**
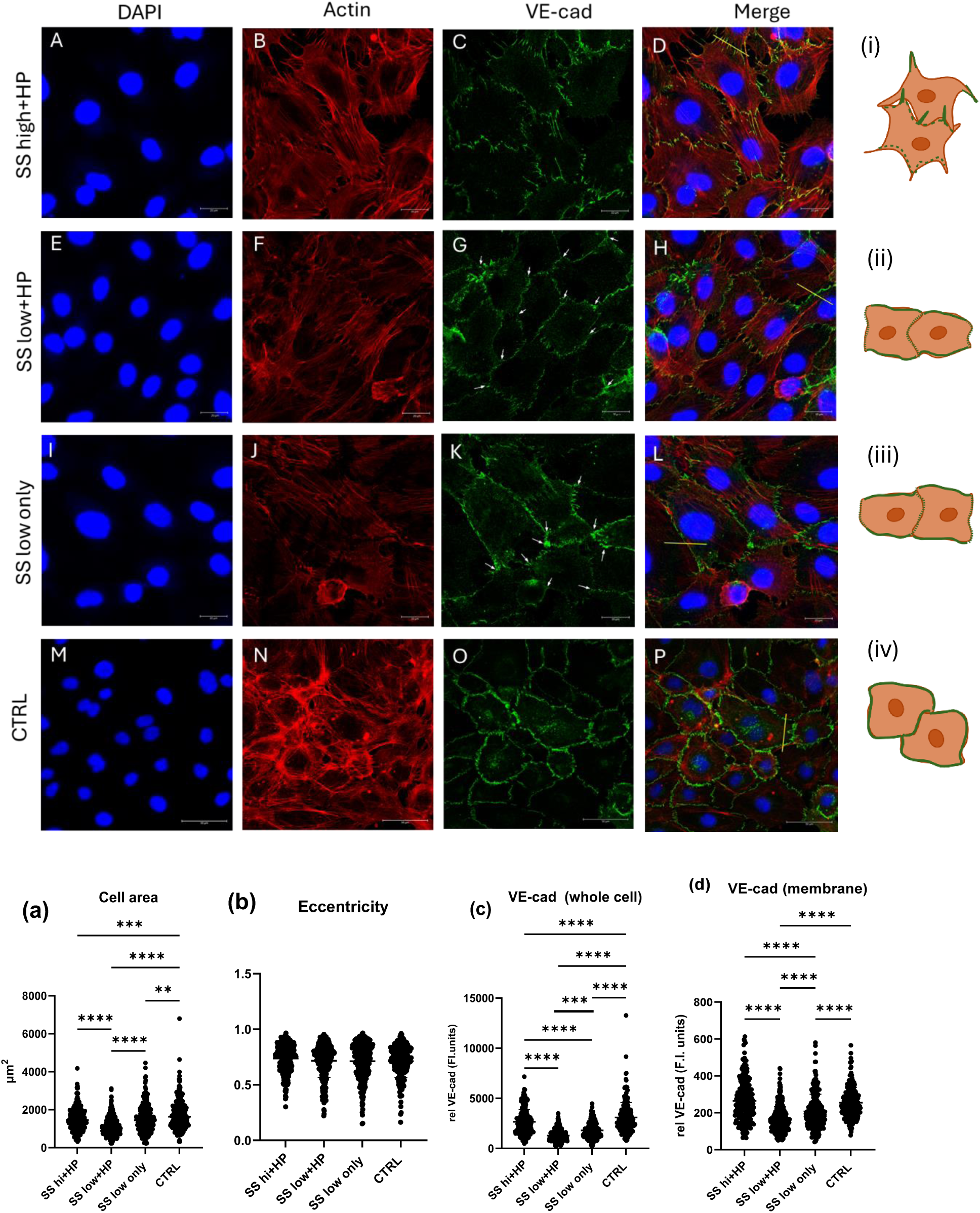
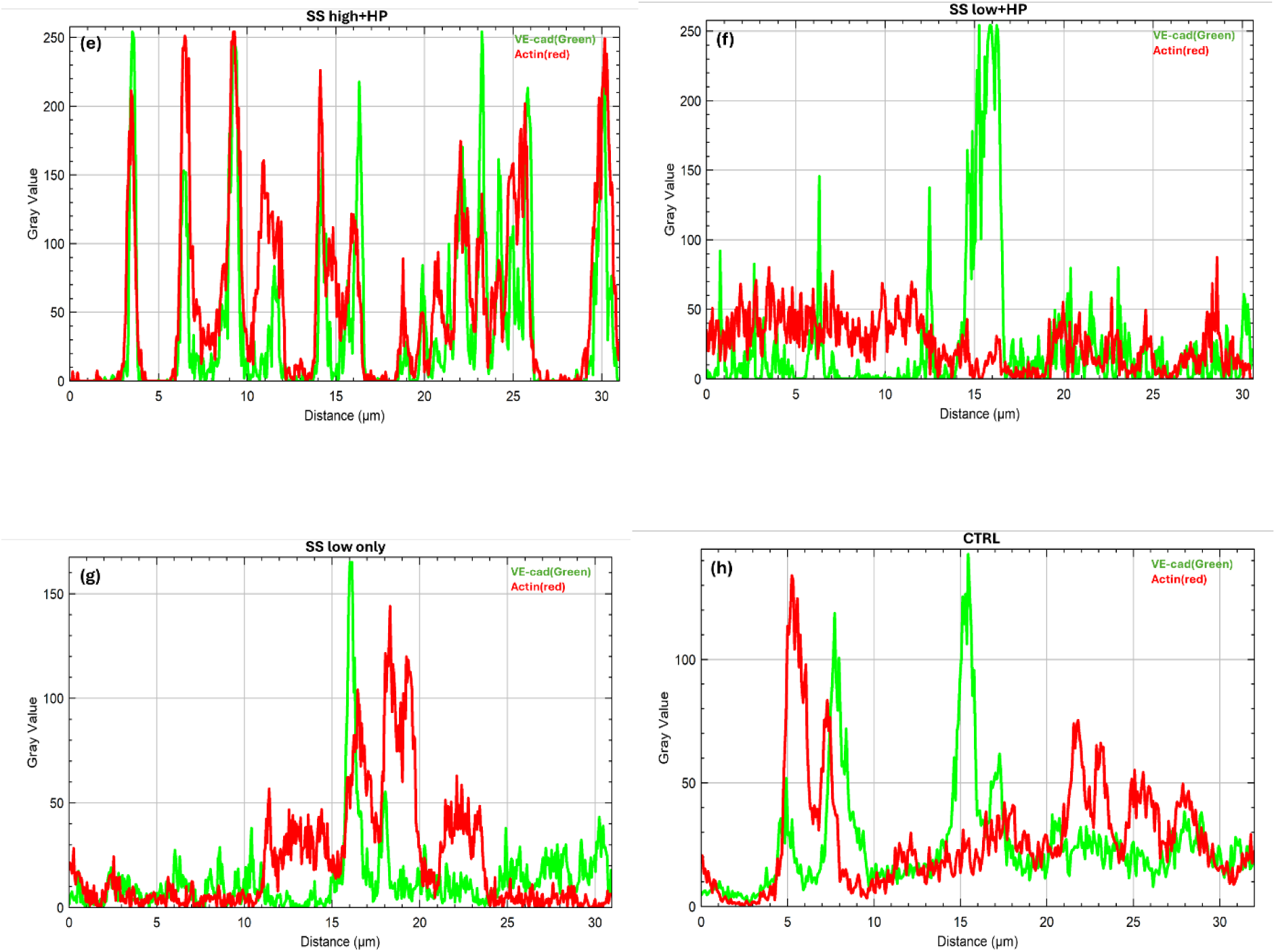
Elevated hydrostatic pressure exposure for 1h at high shear stress causes VE-cadherin finger like protrusions at cell-cell junctions. (A-D) Cells exposed to SS high+HP exhibited serrated, finger like VE-cadherin expression pattern at the cell-cell junctions that co-localised with actin protrusions engulfing into neighbouring cells. In contrast, cells exposed to SS low+HP conditions (E-H) demonstrated continuous VE-cadherin patterning at cell-cell junctions, especially on the sides, whereas the front and back of cells expressed relatively smaller serrated VE-cadherin patterns. Cells exposed to SS low only (I-L) demonstrated VE-cadherin and actin expression pattern similar to that of SS low+HP. Control cells (M-P) demonstrated typical endothelial static cell phenotype with VE-cadherin localised at the cell membrane and thick actin bundles concentrated at the cell cortex. (a) average cell area (b) cell eccentricity (c) VE-cadherin relative fluorescence intensity in whole cell (d) VE-cadherin relative fluorescence intensity in the cell membrane. (i-iv) Schematic representation of cell morphologies observed in different conditions. Green borders at the cell membrane in the schematic represents VE-cadherin. (e-h) represents the line plots of actin (red) and VE-cadherin (green) for SS high+HP, SS low+HP, SS low only and CTRL based on the yellow line (chosen randomly) drawn in D, H, J and P respectively. N = 3, at least 50 cells were analysed per repeat. Flow direction: Top to bottom. White arrows in G,K: tricellular junctions. Mag = 63x (A-L), 40x (M-P)

We then investigated VE-cadherin phenotypic expression pattern and its co-localisation with actin at the cell-cell junctions in different conditions. As shown in Figure 2 (A-D), cells in SS high+HP conditions demonstrated a serrated VE-cadherin pattern in finger- like projections, which protruded into the neighbouring cells as shown in Figure 3A (white arrowheads). As shown in Figure 2 (e) line plot, there was extensive co- localisation of VE-cadherin and thick actin stress fibres in these serrated junctions that protruded away from the cells i.e. the VE-cadherin fingers were orientated orthogonal to their neighbouring actin fibres (refer Supplementary Figure S2 for orientation analysis).

**Figure 3.**
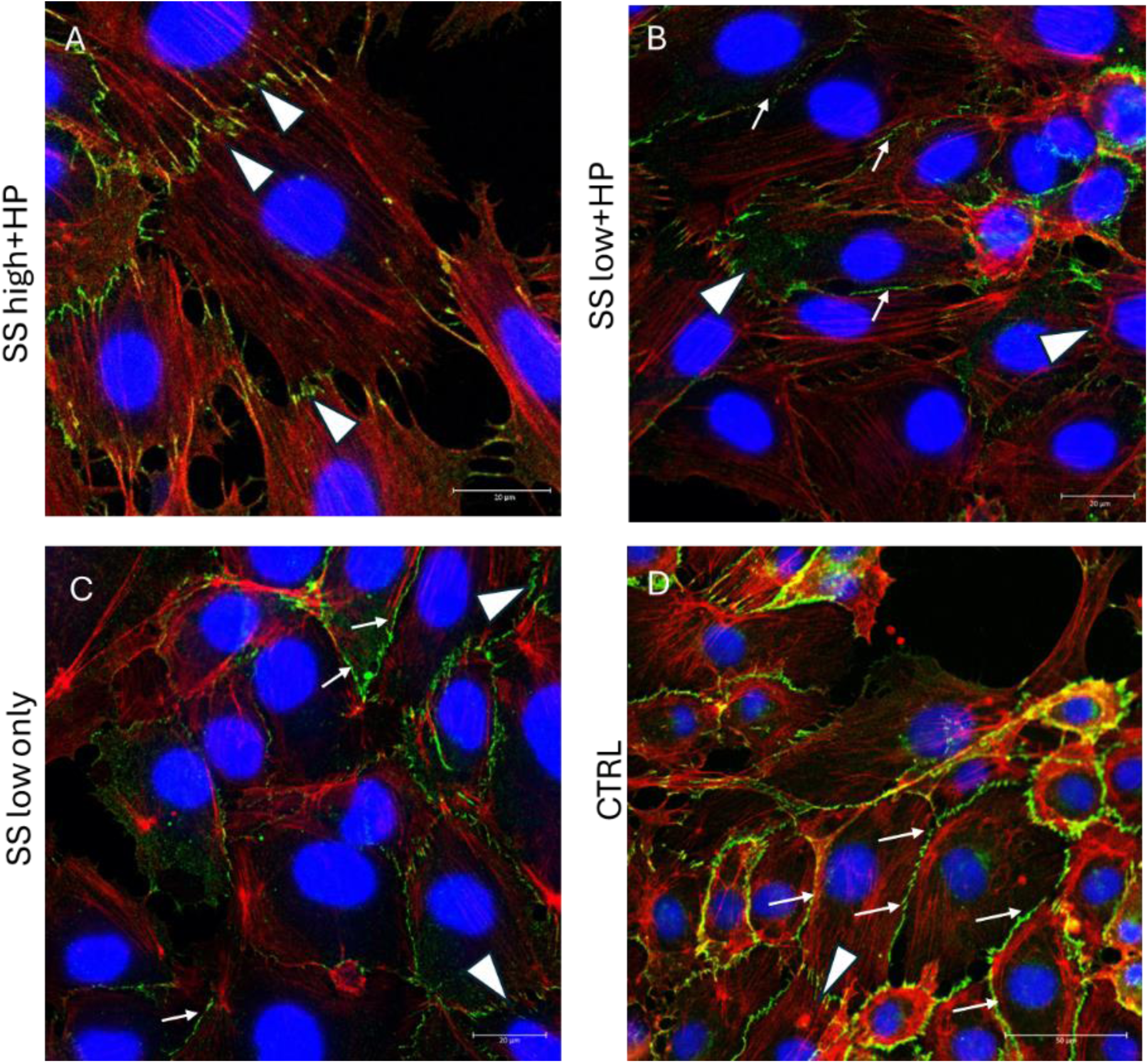
Junctional remodelling in different flow conditions. (A) SS high+HP (B) SS low+HP (C) SS low only and (D) static CTRL. White arrows indicate continuous VE-cadherin junctions, white arrow heads indicate serrated VE-cadherin fingers. N = 3, at least 50 cells were analysed per repeat. Flow direction: Top to bottom. Mag = 63x (A-C), 40x (D)

The lack of continuous, linear VE-cadherin junctions in the SS high+HP conditions indicated that the cells are predominantly in a remodelling phase in which VE- cadherin/actin finger like protrusions produce a pushing or pulling force ^33, 34^ on the junctions. Such VE-cadherin/actin rich protrusions at remodelling cell-cell junctions have been referred to by several studies under different names such as zipper-like adherens junctions ^34^, punctuate adherens junctions ^35^ and serrated adherens junctions. For the rest of this article, we will to refer to these VE-cadherin-actin rich protrusions as VE-cadherin fingers, a term previously employed by Hayer et al. (2016)^36^.

In contrast to the SS high+HP conditions, cells in SS low+HP exhibited both smooth, continuous VE-cadherin junctions mostly at the sides of the cells as shown in Figure 3 (B) (white arrows) whereas the front and back exhibited relatively smaller VE-cadherin serrated patterns as shown in Figure 3B (white arrowheads). Cells with continuous junctions consisted of cortical actin fibres that did not co-localise with VE-cadherin, as shown in Figure 2 (f), which indicates a stable, mature adherens junction as reported by previous studies ^37, 38^. There was an increased number of cells with tri-cellular junctions, as shown in Figure 2G (white arrows), compared to the SS high+HP conditions, another marker of stable junctions and non-motile cell phenotype. Cells in the SS low only conditions exhibited similar tri-cellular junctions, as shown in Figure 2K (white arrows), with a few serrated VE-cadherin junctions as shown in Figure 3C (white arrowheads).

These results indicate that the mechanotransduction from elevated hydrostatic pressure is moderate at low shear stresses (indicated by the presence of continuous VE-cadherin junctions) but can have a significant effect at higher shear stresses (indicated by the presence of distinct VE-cadherin fingers at junctions)

Since the pressure drop decreases across the channel length, as shown in Figure 1 (C,D), endothelial cells closer to the inlet (high-pressure regions relative to the outlet) exhibited characteristics different to that of cells near the outlet (low-pressure regions) Cells closer to the inlet demonstrated increased cell density compared to the cells near the outlet (refer Supplementary Table T1). Further, SS high+HP cells in the sparsely populated regions demonstrated multi-filopodial protrusions (refer Supplementary S3) and reduced VE-cadherin finger formation compared with the cells in the confluent regions which formed prominent VE-cadherin fingers that engulfed deep into the neighbouring cells (refer Supplementary S4), thus demonstrating the significant influence of cell density on VE-cadherin patterning in the SS high+HP conditions.

### 3.2 Piezo-1 inhibition leads to increased VE-cadherin localisation at cell-cell junction

Since piezo-1 channels are known to be extremely sensitive to any mechanical force experienced by the endothelial membrane lipid bilayer ^39, 40^, we next investigated the impact of pharmacologically blocking piezo-1 on VE-cadherin expression pattern at elevated pressure. Cells were exposed to all three flow regimes in the presence of 5 µM GsMTx4 which is known to selectively inhibit piezo-1 channels at micromolar concentrations ^29^.

Blocking piezo-1 in the SS high+HP conditions led to a significant reduction in the average cell area compared to the non-inhibited conditions as shown in Figure 4 (a) and most cells developed thick cortical actin mesh (all conditions were significant compared to the control in Figure 4 (a)). Interestingly, there was a significant increase in the VE-cadherin thickness at the cell membrane but a reduction in the VE-cadherin finger length and filopodia protrusion length compared to the non-inhibited conditions as shown in Figure 4 (e,f,g) respectively. In comparison, blocking piezo-1 in the SS low+HP conditions showed no significant changes in the average cell area and filopodial lengths but a significant increase in the VE-cadherin membrane thickness (similar to SS high+HP). Cells predominantly exhibiting radial, cortical actin fibres and an increase in membrane VE-cadherin F.I. compared to SS high+HP was observed as shown in Figure 5(d).

**Figure 4.**
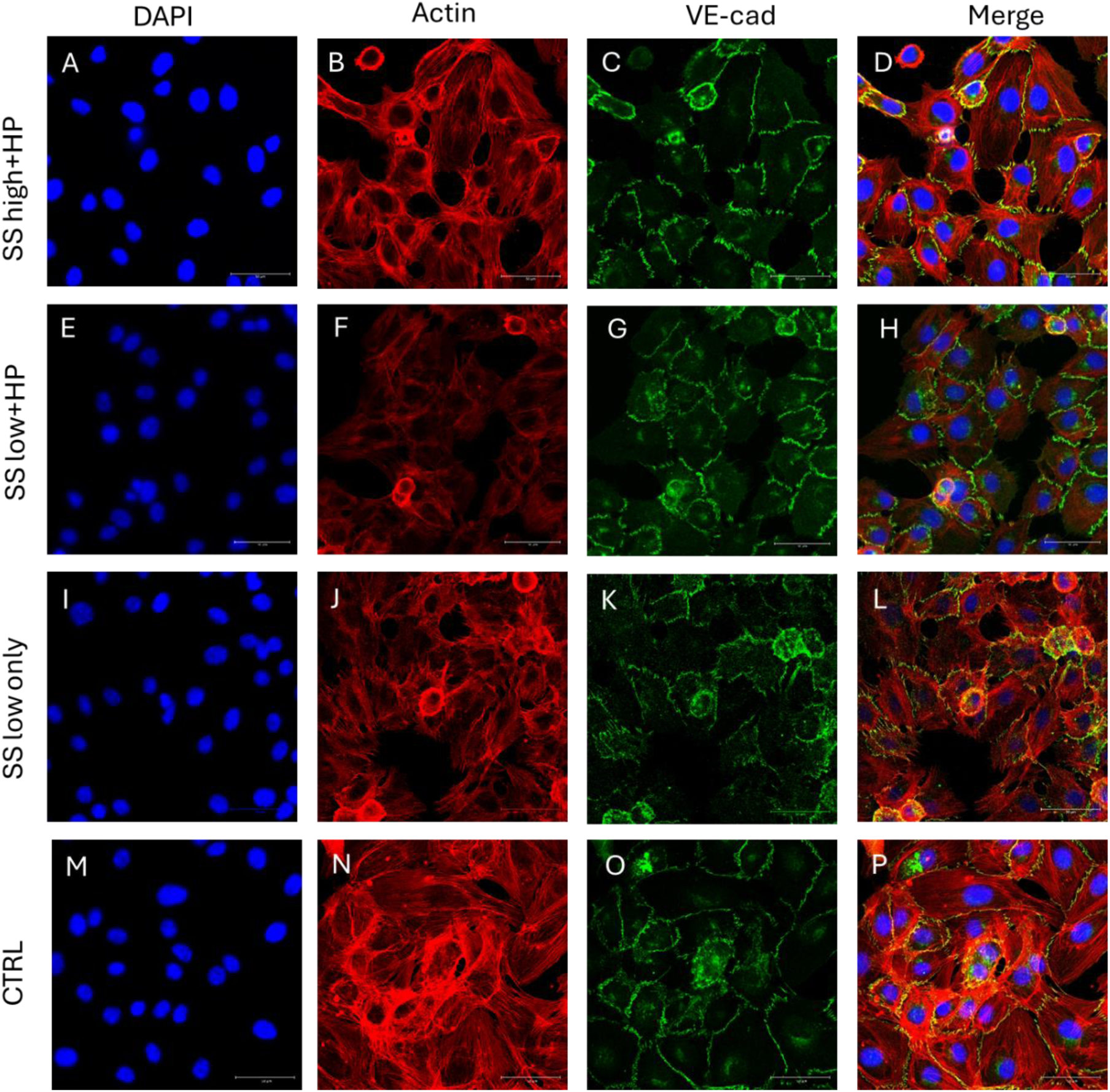
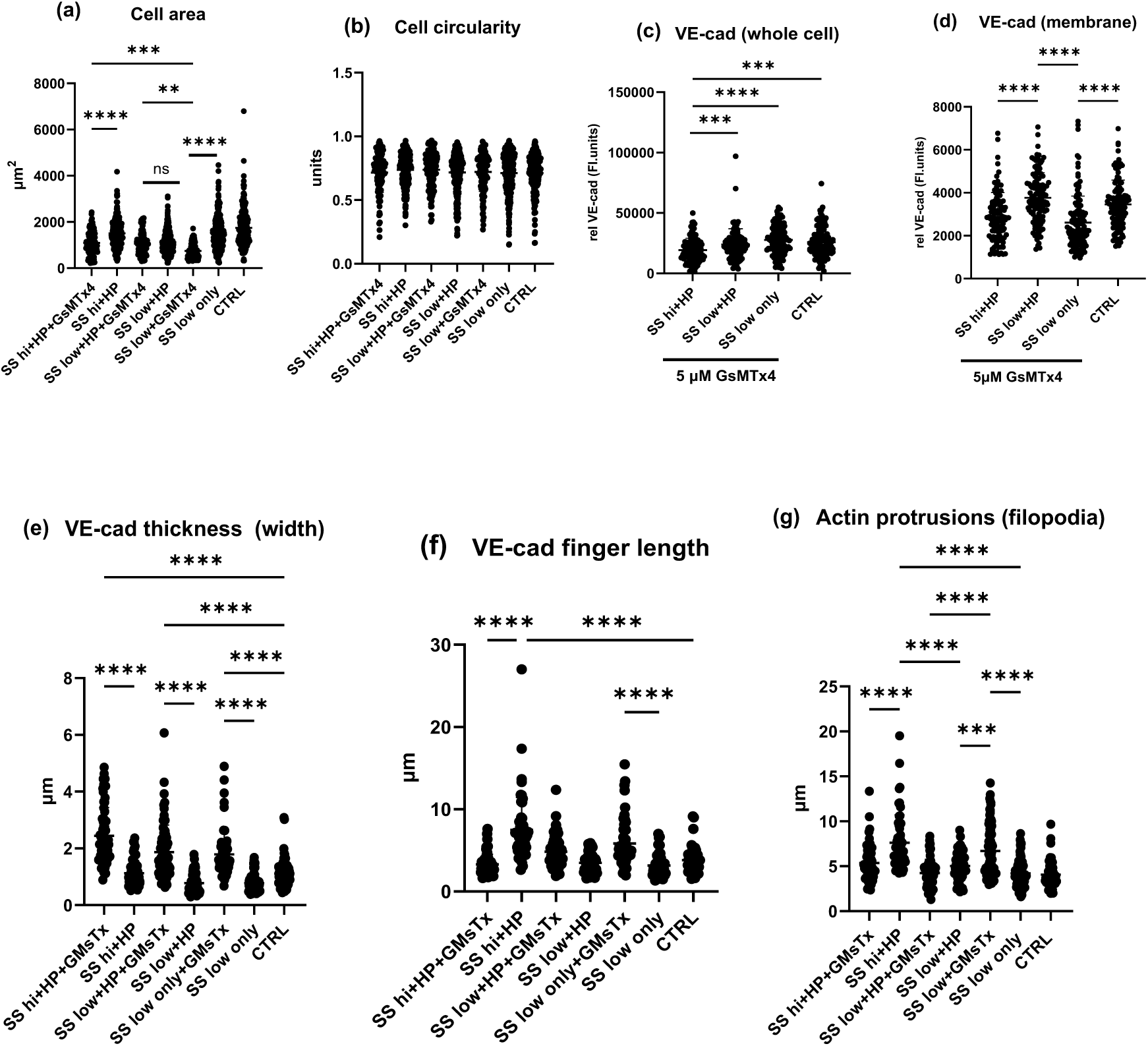
Influence of piezo-1 inhibitor on the effect of elevated hydrostatic pressure. (A-D) SS high+HP demonstrated thick, continuous, stable VE-cadherin junctions at the cell membrane although the number and length of VE-cadherin fingers decreased compared to the SS high+HP conditions. (E-H) SS low+HP demonstrated similar thick VE-cadherin lining at the cell-cell junctions although they covered more cell-cell boundaries compared to the SS high+HP. (I-L) SS low only condition also demonstrated increased number and thickness of VE-cadherin fingers compared to both the pressure conditions in the presence of GsMTx4. (M-P) CTRL cells demonstrated stable, mature junctions consisting mostly of continuous VE-cadherin expression at the cell membrane. (a) average cell area (b) cell circularity (c) VE-cadherin relative F.I. in whole cells (d) VE-cadherin relative F.I. in the cell membrane (e) average VE-cadherin thickness (f) average VE-cadherin finger length (g) average filopodial protrusion length. N = 3, at least 50 cells were analysed per repeat. Flow direction: Top to bottom. Mag = 40x

**Figure 5.**
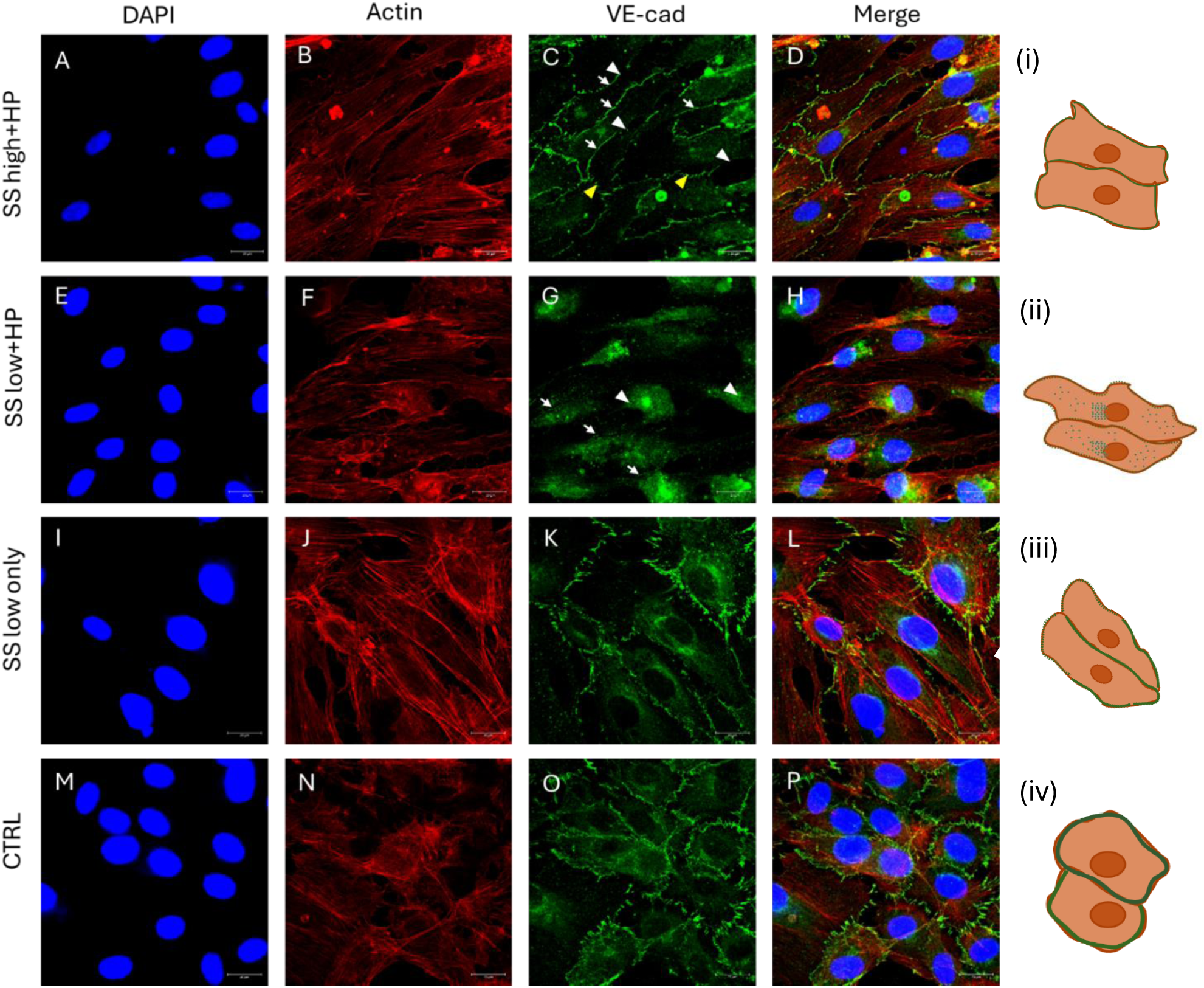
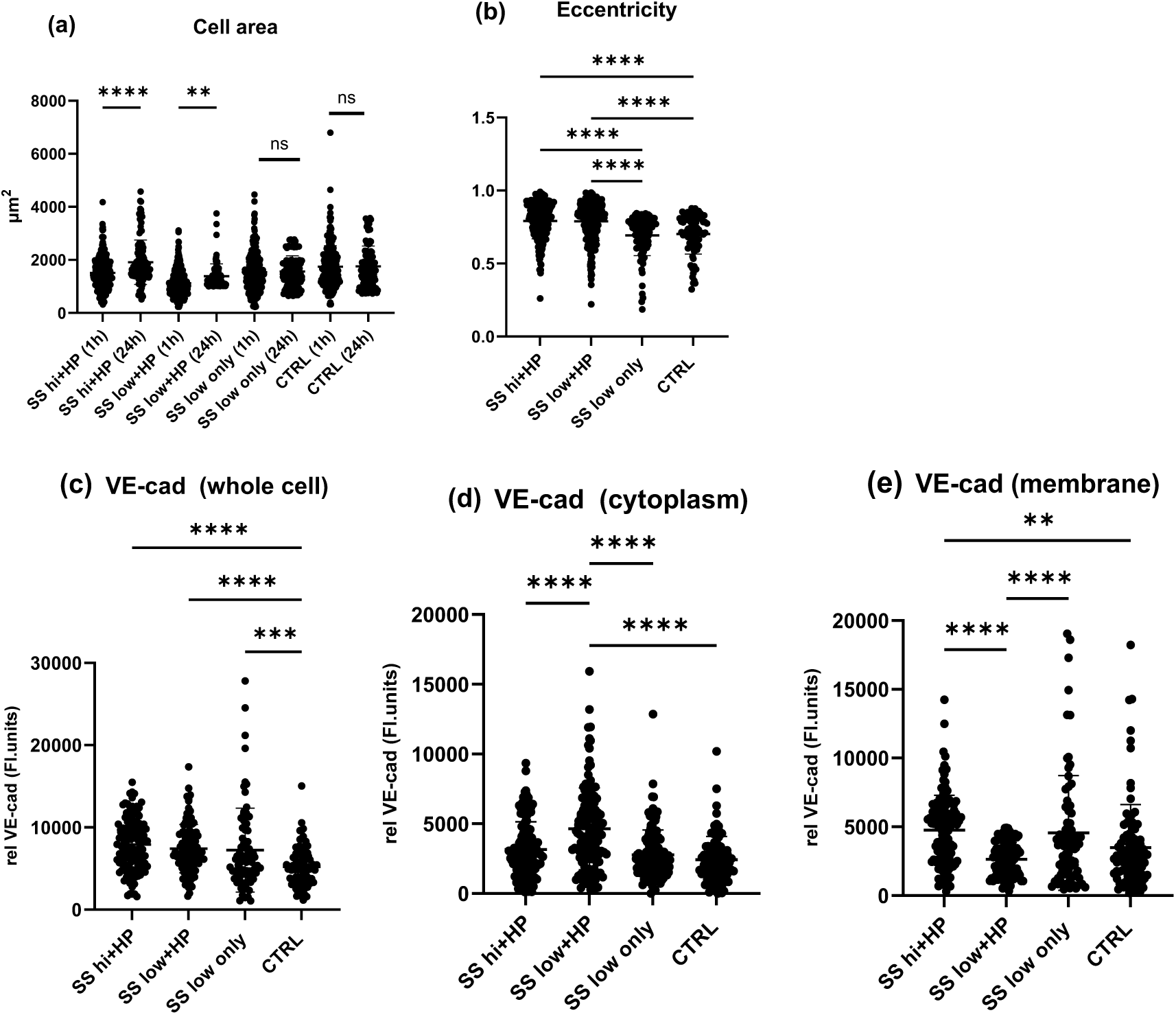
Influence of long-term (24h) exposure to elevated hydrostatic pressure on VE-cadherin expression. (A-D) SS high+HP, (E-H) SS low+HP, (I-L) SS only, (M-P) static control cells. Although there was no difference in relative VE-cadherin F.I. when analysing whole cells, there was a marked difference in the cytoplasmic and membrane regions with SS low+HP showing higher VE-cadherin expression in the cytoplasm compared to the membrane indicating internalisation. (a) average cell area in different flow conditions after 1h and 24h exposure (b) cell eccentricity after 24h exposure (c) VE-cadherin relative F.I. in whole cell (d) VE-cadherin relative F.I. in the cell cytoplasm (e) VE-cadherin relative F.I. in the cell membrane (i-iv) Schematic representation of cell morphologies observed in different conditions. Green borders at the cell membrane in the schematic represent VE-cadherin. N = 3, at least 50 cells were analysed per repeat. Flow direction: Left to right. Mag = 63x

Surprisingly, inhibiting piezo-1 in the SS low only conditions demonstrated a significant increase in the VE-cadherin membrane thickness but also in the VE-cadherin finger length and filopodial length, which was comparable to that of SS high+HP conditions. This observation was interesting as it demonstrated that inhibiting piezo-1 during the application of a hydrostatic pressure head can alter VE-cadherin finger patterning differently: it can either lead to decreased VE-cadherin finger length as observed in SS high+HP or it does not alter the VE-cadherin finger length as observed in SS low+HP, whereas the removal of hydrostatic pressure in SS low only conditions caused an increase in the VE-cadherin finger length. Further, since Piezo-1 channels sense mechanical perturbations across cell membrane by tethering to actin via cadherin/β- actin complexes ^40^, we analysed how inhibiting piezo-1 impacted the actin cytoskeletal arrangements. We observed that inhibition of piezo-1 led to a reduction in filopodia formation at cell edges (line plots quantification in supplementary Figure S5), instead promoting radial actin mesh arrangement as shown in Figure 4 (B,F,J). This suggests that piezo-1 plays a key role in the mechanotransduction of hydrostatic pressure to regulate VE-cadherin/actin protrusions at cell-cell junctions.

### 3.3 Prolonged exposure to elevated hydrostatic pressure disrupts junctional VE-cadherin but only at low shear stress

We next investigated the long-term effects of elevated hydrostatic pressure application at both shear stresses. Cells exposed to SS high+HP for 24 h demonstrated an elongated morphology, as shown in Figure 5 (A-D), with an increased average cell area and significantly less circularity compared to the control cells, as shown in Figure 5 (M-P) and quantified in Figure 5 (a,b). Cells also demonstrated aligned actin stress fibres covering the entire cell body and forming extensive connections with the neighbouring cells with increased actin/VE-cadherin co-localisation at the membrane (line plots quantification in supplementary Figure S6). VE-cadherin clustered predominantly at the membranes as continuous junctions (Figure 5C, white arrows) with intermittent disruptions (Figure 5C, white arrowheads) and some cells exhibiting immature VE-cadherin fingers (Figure 5C, yellow arrowheads).

In comparison, cells exposed to SS low+HP for 24 h demonstrated almost a substantial disruption of VE-cadherin at the membrane with only a faint, thin VE-cadherin lining at cell-cell contacts (Fig 5G, white arrows). Interestingly, there was an increased localisation of VE-cadherin in the cell cytoplasm (Fig 5G, white arrowheads), and the same has been quantified in Figure 5 (d,e). This suggests that prolonged exposure to elevated hydrostatic pressure under low shear stress caused endocytosis of VE-cadherin. It is also interesting to note the markedly different impact of 24 h pressure exposure compared to the 1 h pressure exposure regimes on the cell morphologies, which is summarised below in Table 2:

**Table 2.** Description of cell morphological characteristics in response to different flow conditions.

|  | **SS low+HP (1h)** | **SS low+HP (24h)** | **SS high+HP (1h)** | **SS high+HP (24h)** |
| --- | --- | --- | --- | --- |
| **Cell morphology** | Polygonal | Elongated | Polygonal | Elongated |
| **VE-cadherin pattern** | Continuous VE-cadherin junctions | Substantial disruption of VE-cadherin junctions | Engulfing VE-cadherin fingers | Continuous VE-cadherin junctions (intermittently disrupted) |
| **VE-cadherin fingers** | Short VE-cadherin with reduced actin/VE-cadherin co-localisation | No VE-cadherin fingers | Long, prominent VE-cadherin fingers with increased actin/VE-cadherin co-localisation | Short VE-cadherin with reduced actin/VE-cadherin co-localisation |
| **Cytoplasmic VE-cadherin** | Minimal cytoplasmic VE-cadherin | Elevated cytoplasmic VE-cadherin localisation | Minimal cytoplasmic VE-cadherin | Minimal cytoplasmic VE-cadherin |
| **Bi-tri-cellular VE-cadherin junctions** | Increased bi-tri-cellular junctions | No bi-tri-cellular junctions | No bi-tri-cellular junctions | Presence of bi-tri-cellular junctions |
| **Filopodial protrusions** | No significant filapodial protrusions at cell edges | Increased filapodial protrusions at cell edges | Increased filapodial protrusions at cell edges | Reduced filapodial protrusions at cell edges |
| **Actin cytoskeleton arrangement** | Radial actin stress fibres arrangement | Parallely aligned stress fibres | Stress fibres forming protrusions at cell edges | Parallely aligned stress fibres |

### 3.4 VE-cadherin endocytosis is reversible upon inhibiting piezo-1

Similar to the short-term exposure conditions, we next investigated if the VE-cadherin endocytosis observed in the SS low+HP 24 h exposure condition was influenced by piezo-1 mechanotransduction. For these experiments, the cells exposed to SS low+HP for 24 h were then post-treated to a growth medium containing 5 µM GsMTx4 for either 4 h or 12 h, after which the cells were fixed for immunostaining. As can be noticed from Figure 6 (A-C), treating cells to 5µM GsMTx4 for 4 h led to partial reassembling of VE-cadherin at cell-cell junctions, whereas 12 h treatment to 5 µM GsMTx4 led to formation of stable VE-cadherin junctions along the cell-cell contacts.

**Figure 6.**
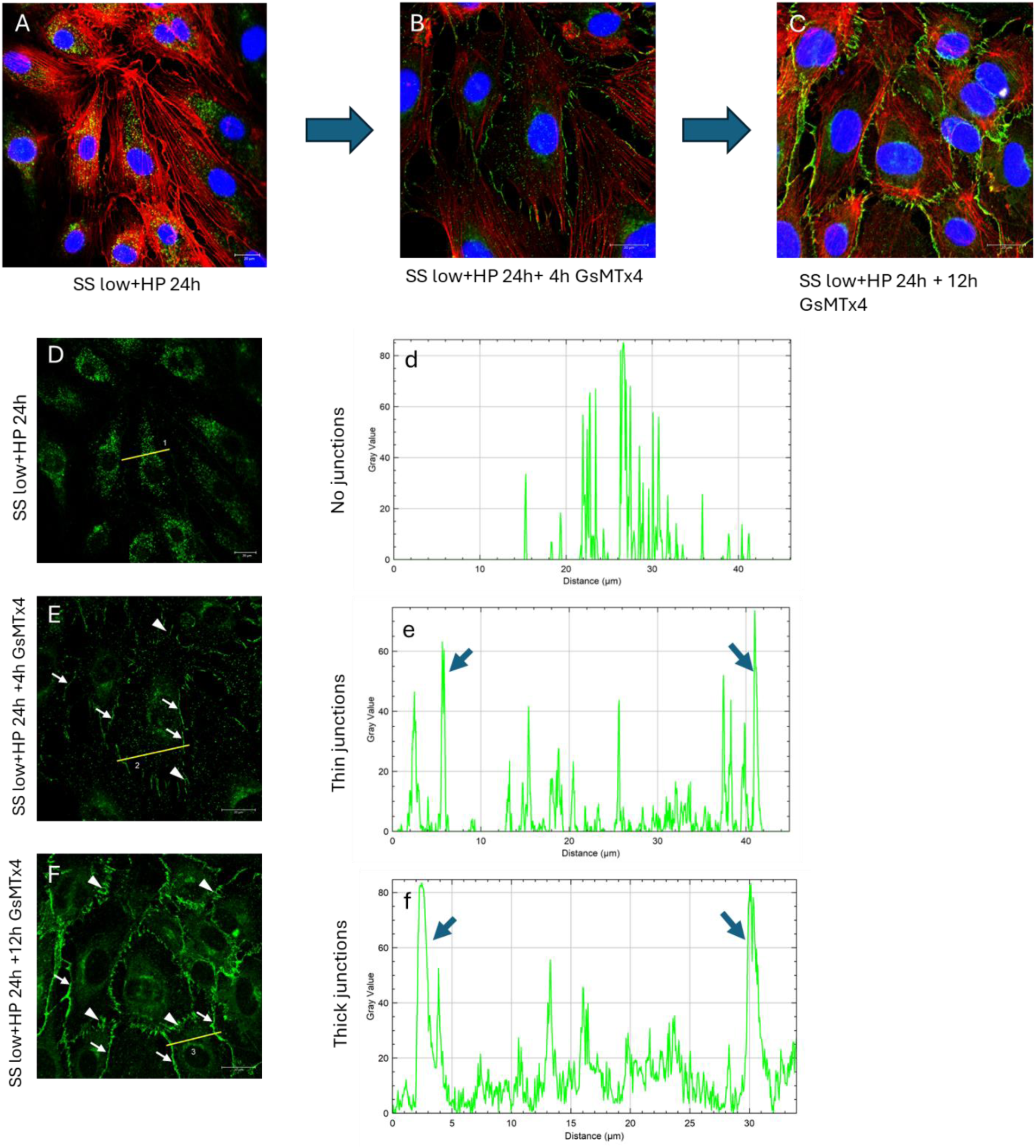
Piezo-1 dependent reversible nature of VE-cadherin endocytosis. **(A)** Cells exposed to SS low+HP for 24h (B) Cells exposed to SS low+HP for 24h followed by treatment to 5µM GsMTx4 for 4h and (C) Cells exposed to SS low+HP for 24h followed by treatment to 5µM GsMTx4 for 12h. It can be observed that cells treated to GsMTx4 for 4h post SS low+HP exposure demonstrated reduced VE-cadherin cytoplasmic endosomes and exhibited development of thin, continuous VE-cadherin junctions (denoted by the white arrows in (E) and blue arrows in (e)). Upon treating the cells to 12h of GsMTx4 post SS low+HP exposure, the cells demonstrated the formation of thick, continuous junctions (denoted by the white arrows in (F) and blue arrows in (f)) as well as immature, VE-cadherin fingers (denoted by the white arrowheads in (F)). Flow direction: Top to bottom. Mag = 40x (A,D), 63x (B,C,E,F)

Further, the VE-cadherin line plots in Figure 6 (d,e,f) demonstrate the gradual formation of continuous VE-cadherin junctions with increasing GsMTx4 post-treatment time.

Figure 6 (e) demonstrates the formation of thin VE-cadherin junctions (white arrows in Figure 6 (E)) and decreased cytoplasmic VE-cadherin endosomes after 4 h GsMTx4 post-treatment compared to Figure 6 (d). After 12 h GsMTx4 post-treatment, thick, continuous VE-cadherin junctions (white arrows in Figure 6 (F)) with no significant cytoplasmic VE-cadherin endosomes were observed as shown in Figure 6 (f). Further, immature, short VE-cadherin fingers were also observed after the 12h GsMTx4 post- treatment (white arrowheads in Figure 6 (F)). These results demonstrate the crucial mechanotransduction role of piezo-1 in regulating VE-cadherin endocytosis dynamics.

## 4 Discussion

The pathophysiology of several diseases, such as ACS, involves increased pressure experienced by the vasculature at a ‘low-flow to no-flow’ condition, which causes microvascular dysfunction and tissue degeneration ^41, 42^. In comparison to shear stress, the mechanobiological role of elevated hydrostatic pressure in the progression of vascular dysfunction has surprisingly received limited attention ^43, 44^. In the current study, using a microfluidic system, we investigated the combined influence of low shear stresses and elevated hydrostatic pressure on endothelial cell behaviour.

Our study demonstrated that without the addition of any inflammatory stimuli, a 1 h exposure to elevated hydrostatic pressure at high but not low shear stress led to VE-cadherin finger protrusions at the cell-cell junctions. Previously, Hayer et al. (2016) reported that such VE-cadherin finger like structures are double plasma membrane bound tubular projections that involve VE-cadherin/actin rich protrusions from the leader cells being engulfed by the plasma membrane of the follower cells, thus creating a polarity in cells and generating actomyosin contractility to promote migration ^36^. More interestingly, our study showed that despite demonstrating similar cell confluency, short term exposure to SS high+HP resulted in VE-cadherin finger formation whereas SS low+HP demonstrated continuous VE-cadherin junctions. This observation was contrary to past studies that showed that serrated VE-cadherin junctions to be a marked feature of sub-confluent cell density and continuous VE-cadherin junctions are characteristic of confluent cell density ^45, 46^. A possible explanation for this cell density independent variable VE-cadherin patterning could be that the cells produce the same level of endogenous VE-cadherin irrespective of the cell density (as a part of the cell energy conservation mechanism to avoid the constant synthesis of new proteins ^45^) as previously stated ^47^ . The type and magnitude of mechanical stimuli would then determine the VE-cadherin patterns i.e. continuous, stable junctions versus finger like projections at the junction. It is also important to note that the effect of hydrostatic pressure is temporally variable. While SS low+HP at 1h formed stable bi-cellular and tri- cellular junctions, SS low+HP at 24h demonstrated VE-cadherin internalisation. This demonstrates that the impact of hydrostatic pressure can have a protective, neutral or disruptive effect depending upon not just the shear stress magnitude but also the exposure time which agrees with previous reports ^16, 48^. In an interesting study highlighting the variable temporal effect of hydrostatic pressure, Prystopiuk (2018) reported a two-phase response of endothelial cells to hydrostatic pressure wherein an acute exposure of 1 h to 100 mmHg did not impact VE-cadherin distribution whereas a chronic exposure of 24 h disrupted VE-cadherin at the membrane which is similar that observed in the current study ^49^.

Though several candidates such as Ca^2+^channels ^50^, TRPV1 ^51^, aquaporins ^48^ have been proposed as mechanical sensors of pressure changes, the exact mechanism by which transient changes in hydrostatic pressure are sensed by endothelial cells still eludes cell biologists. Piezo-1 is a transmembrane homotrimer protein chiefly responsible for mechanical force sensing and regulating blood pressure in vascular endothelial cells ^52^. In the current study, short term inhibition of piezo-1 lead to increased accumulation of VE-cadherin at cell-cell contacts in both the elevated pressure conditions. This observation agrees with previous studies wherein elevated hydrostatic pressure caused disruption of VE-cadherin at junctions in a piezo-1 dependent manner and caused hyperpermeability ^53, 54^. However, an exact opposite effect was reported by Zhong et al. wherein piezo-1 activation protected cells from VE-cadherin internalisation and promoted junctional integrity in response to mechanical stretching ^55^. These studies suggest that different mechanical stimuli can impact the junctional integrity differently, with piezo-1 being barrier disruptive in response to elevated hydrostatic pressure but being barrier protective in response to cyclic mechanical stretch.

Although past studies have shown VEGF ^56^, histamine ^57^ and TNF-α ^58^ treatments to cause VE-cadherin internalisation, no study has, to our knowledge, reported on hydrostatic pressure (in the absence of any chemical stimuli) induced VE-cadherin internalisation. Several mechanisms of VE-cadherin internalisation have been reported. Xiao et al. (2005) demonstrated that the internalisation of endogenous VE-cadherin is mediated via clathrin but not in a caveolae-dependent manner ^59^. Another study by Gavard and Gutkind (2006) demonstrated a mechanism that involves β-arrestin mediated endocytosis of VE-cadherin in clathrin coated vesicles ^60^. Further investigations are required to shed light on the exact molecular mechanisms of VE-cadherin internalisation since past studies have shown both clathrin ^61^ or caveolae mediated ^62^ VE-cadherin internalisation ^63^. Further, following internalisation, VE-cadherin can either be recycled back to the cell membrane or degraded via the lysosomal machinery ^64^. In the current study, inhibiting piezo-1 led to re-clustering of VE-cadherin at cell-cell contacts. This observation is in agreement with one past in vivo study which showed that increased vascular pressure post stenotic constriction reduced the expression of VE-cadherin (no internalisation was reported) and that this reduction in VE-cadherin levels was abrogated upon deleting piezo-1 gene or blocking piezo-1 using GsMTx4, thus demonstrating the inverse relationship between piezo-1 and VE-cadherin ^65^. Although deciphering the mechanism behind this relationship between VE-cadherin and piezo-1 was beyond the scope of the current study, it would nevertheless be interesting to investigate the spatiotemporal relationship between VE-cadherin, piezo-1 and the recycling of small GTPase Rab11a at elevated hydrostatic pressure to understand the VE-cadherin recycling mechanism further.

## 5 Conclusions

This study reports on the temporal and shear stress dependent effects of elevated hydrostatic pressure on the dynamics of VE-cadherin and actin at the endothelial cell-cell junctions. We show that short-term exposure to high pressure led to the formation of VE-cadherin finger-like projections along actin protrusions at the cell membrane, which was indicative of a remodelling junction suggesting a motile cell phenotype.

Importantly, these VE-cadherin finger projections and actin protrusions were abrogated upon pharmacologically blocking piezo-1 which was a novel observation. We also show that the magnitude of shear stress influences the response of cells to elevated hydrostatic pressure. Long term exposure to low but not high shear stress at elevated hydrostatic pressure led to VE-cadherin internalisation. Finally, we also report that the observed VE-cadherin internalisation was reversible upon blocking piezo-1 in a time dependent manner. Understanding junctional remodelling facilitated by VE-cadherin expression, patterning, internalisation, and recycling in a high-pressure environment has major implications in diseases like ACS, glaucoma, etc., that involve dysregulation endothelial permeability and inflammation. Further investigations into the mechanisms of elevated pressure induced VE-cadherin remodelling at junctions could provide new markers of early diagnosis and novel mechanotherapy interventions for these vascular conditions.

## 7 Data availability

All data supporting the findings of this study are included in the manuscript and its ESI.

## Acknowledgements

This work was supported by funding received from the UK Engineering and Physical Sciences Research Council (EPSRC) Programme Grant PREMIERE (EP/T000414/1). The authors would like to thank Dr. Alexander Brill (University of Birmingham) and Dr. John James (University of Warwick) for their valuable critical discussions and technical suggestions.

## 10 Author contributions

PVB designed, conducted experiments and analysed the data. TA and PPE performed the numerical solutions for flow within a rectangular channel. PVB wrote the manuscript, which was revised by DV, MJHS and LMG. All authors discussed experiments, data analysis and results. All authors approved the final version of the manuscript.

