## Supplementary Data for "Elevated hydrostatic pressure acts via piezo-1 to destabilise VE-cadherin junctions: an endothelium-on-chip study"

### 1.1 Randon alignment of actin fibres in response to 1h exposure to elevated hydrostatic pressure

No particular alignment of actin fibres was seen in response to 1h exposure to elevated hydrostatic pressure across all three conditions.

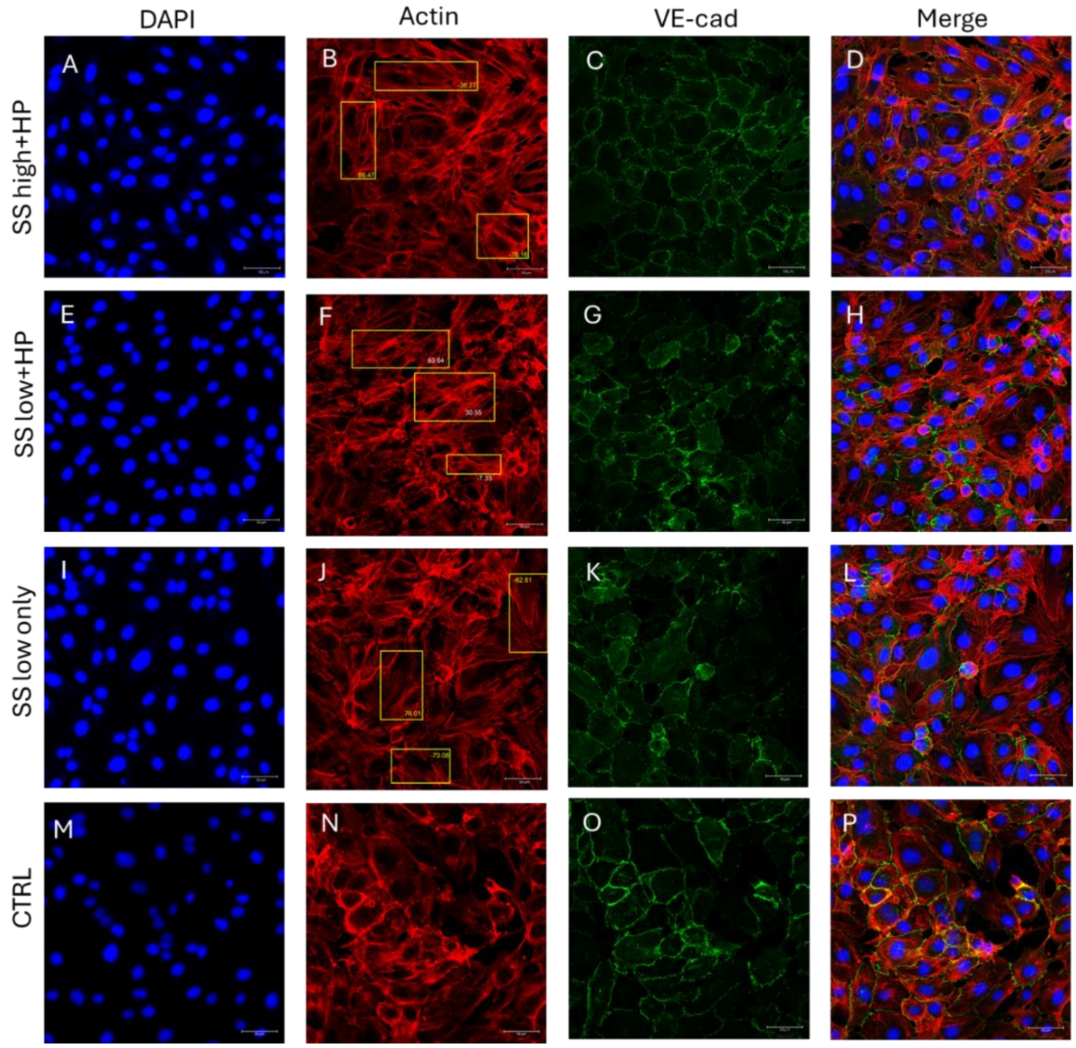


**Figure S1. Short-term (1h) influence of elevated hydrostatic pressure varies depending upon the shear stress magnitude.** A-D) SS high+HP, (E-H) SS low+HP, (I-L) SS only, (M-P) static control cells. Note that in all the flow conditions, irrespective of the hydrostatic pressure or shear stress magnitude, there was no particular alignment of the cells in the direction of the flow (top to bottom) as depicted by the varying orientation angles of actin cytoskeleton in the yellow boxed regions in (B), (F), (J) and (N). Majority of the cells in the SS high+HP conditions demonstrated parallelly aligned actin stress fibres whereas the low shear stress conditions consisted of a mixture of cells exhibiting stress fibres or radial cortical actin structures similar to that seen in the static control cells. N = 3, at least 50 cells were analysed per repeat. Flow direction: top to bottom. Mag = 25x

### 1.2 VE-cadherin finger and actin fibre misalignment at cell-cell junctions

We analysed the orientation angle of VE-cadherin relative to the neighbouring actin fibres which can indicate a remodelling junction in cells in response to a mechanical force. As shown in Figure S2 (A), in the highlighted cell, VE-cadherin fingers and actin aligned in the same direction with peak orientation angles of -70.24° (Figure S2A (i)) and -64.62° (Figure S2A (ii)) respectively. In contrast, in a neighbouring cell highlighted in Figure S2(B), the VE-cadherin fingers orientated at 71.15° ((Figure S2B (i)) while the basal actin fibres demonstrated a radial arrangement with a flattened peak orientation at -52.41° (Figure S2B (ii)). This result agrees with past studies that have shown that in remodelling junctions, VE-cadherin has perpendicular orientation relative to radial actin fibres and that such orientation indicate remodelling junctions [39, 40] .

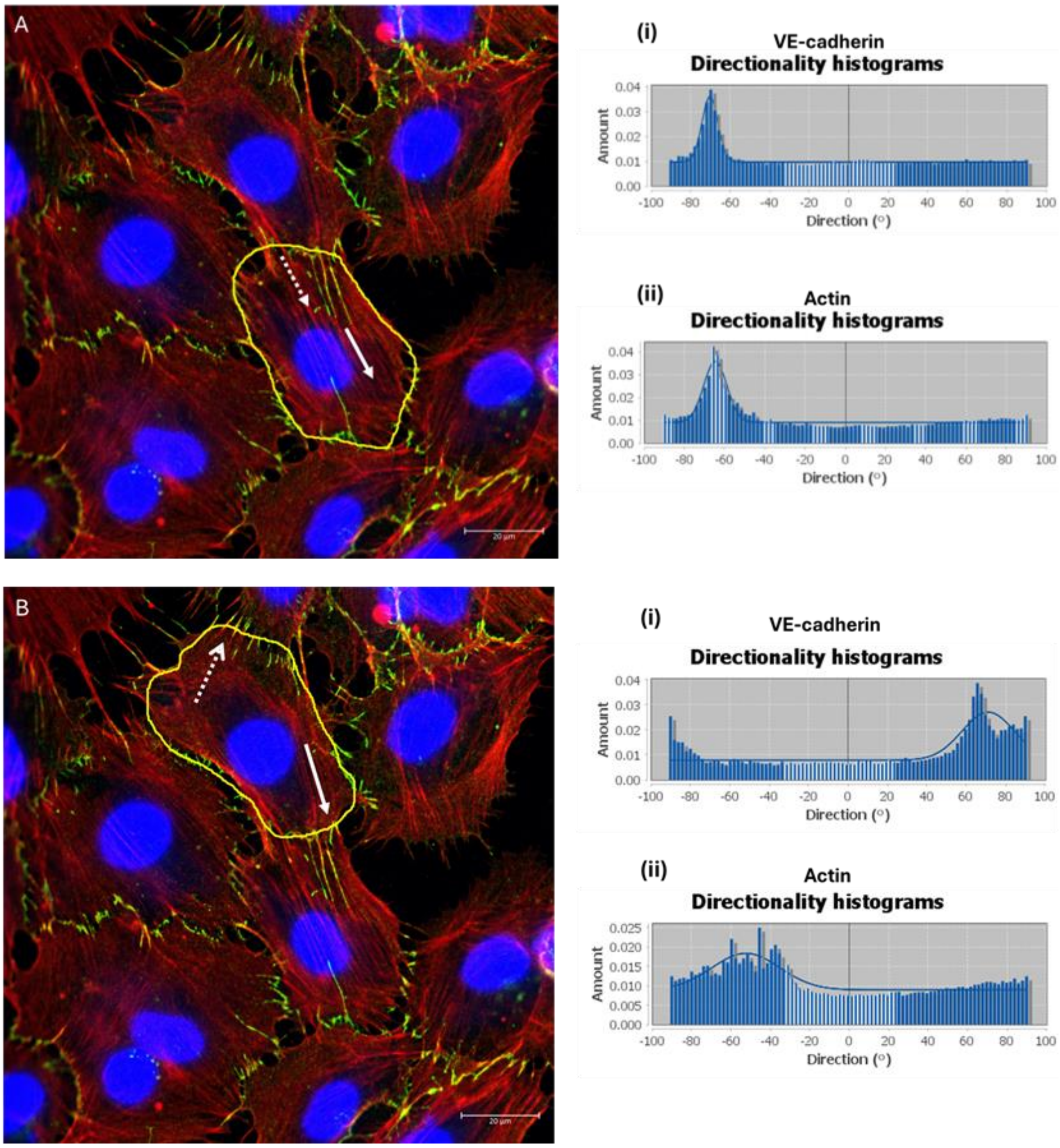


**Figure S2. Mismatch between VE-cadherin fingers and neighbouring actin fibre orientation.** (A) A cell in the SS high+HP condition where the engulfing VE-cadherin fingers (dotted white arrow) aligned with the actin stress fibres (white arrow). As demonstrated by the histogram plots (i) VE-cadherin fingers had a peak orientation angle of -70.24° and actin fibres (ii) had a peak orientation angle of -64.62° (B) A neighbouring cell in the same channel in which the VE-cadherin fingers with a peak orientation angle of 71.15° were protruding away from the dominant direction of radial actin fibres (-52.41°) as demonstrated by the histogram plots. Flow direction: Top to bottom. White arrows- actin fibre orientation, Dotted white arrows- VE-cadherin finger orientation. Mag = 63x

### 1.3 Hydrostatic pressure produces variable effect on cell density across the microfluidic channel

At all three conditions, significantly higher cell numbers were observed in the high-pressure regions (cells closer to the channel inlet) compared to the low-pressure regions (cells closer to the channel outlet) where the cell density was low as shown in Table T1.

**Table T1. Average cell density in the high-pressure (closer to the channel inlet) and low-pressure (closer to the channel outlet) regions of the microfluidic channel. Six 25x images (2580x2580) were analysed per condition.**

| | SS high+HP (mean±SD, n=6) | SS low+HP (mean±SD, n=6) | SS low only (mean±SD, n=6) |
|---|---|---|---|
| High pressure regions | 66.83±1.77 | 85.16±2.26 | 67.66±5.84 |
| Low pressure regions | 45.83±2.40 | 39±3.55 | 34.16±4.48 |

Figure S3 (A,B) represents the cells in the low-pressure regions (3924 Pa) and Figure S3 (D,E) represents the cells in the high-pressure regions (3971 Pa for high shear stress and 3928.4 Pa for low shear stress conditions) for the elevated pressure conditions.

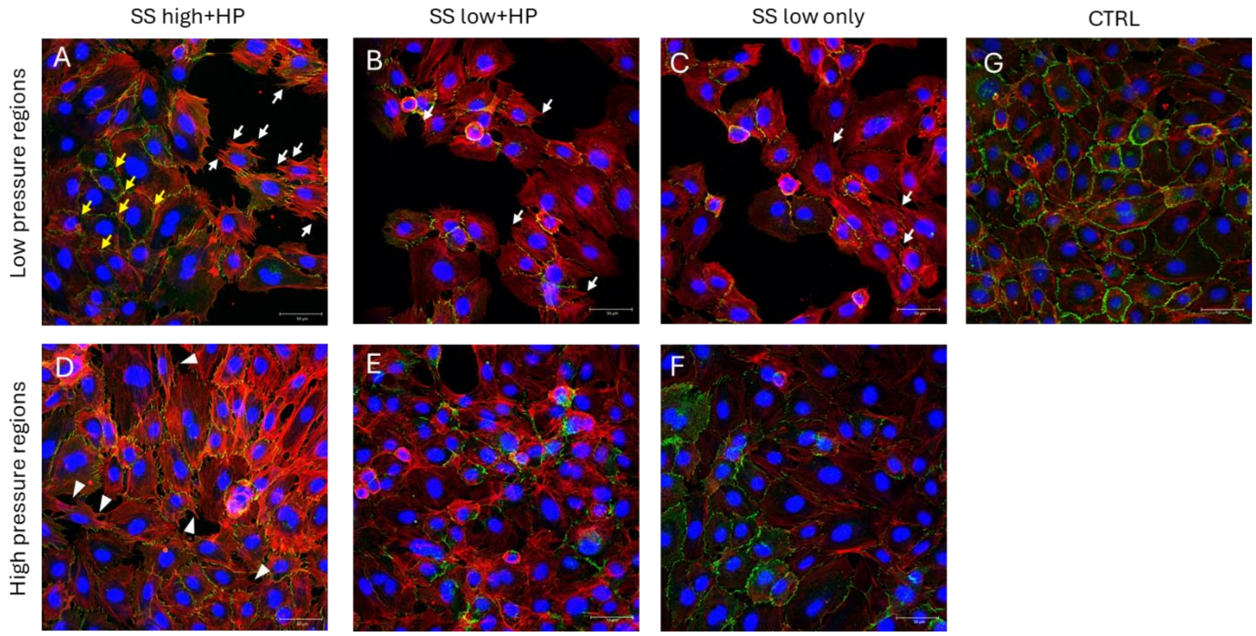

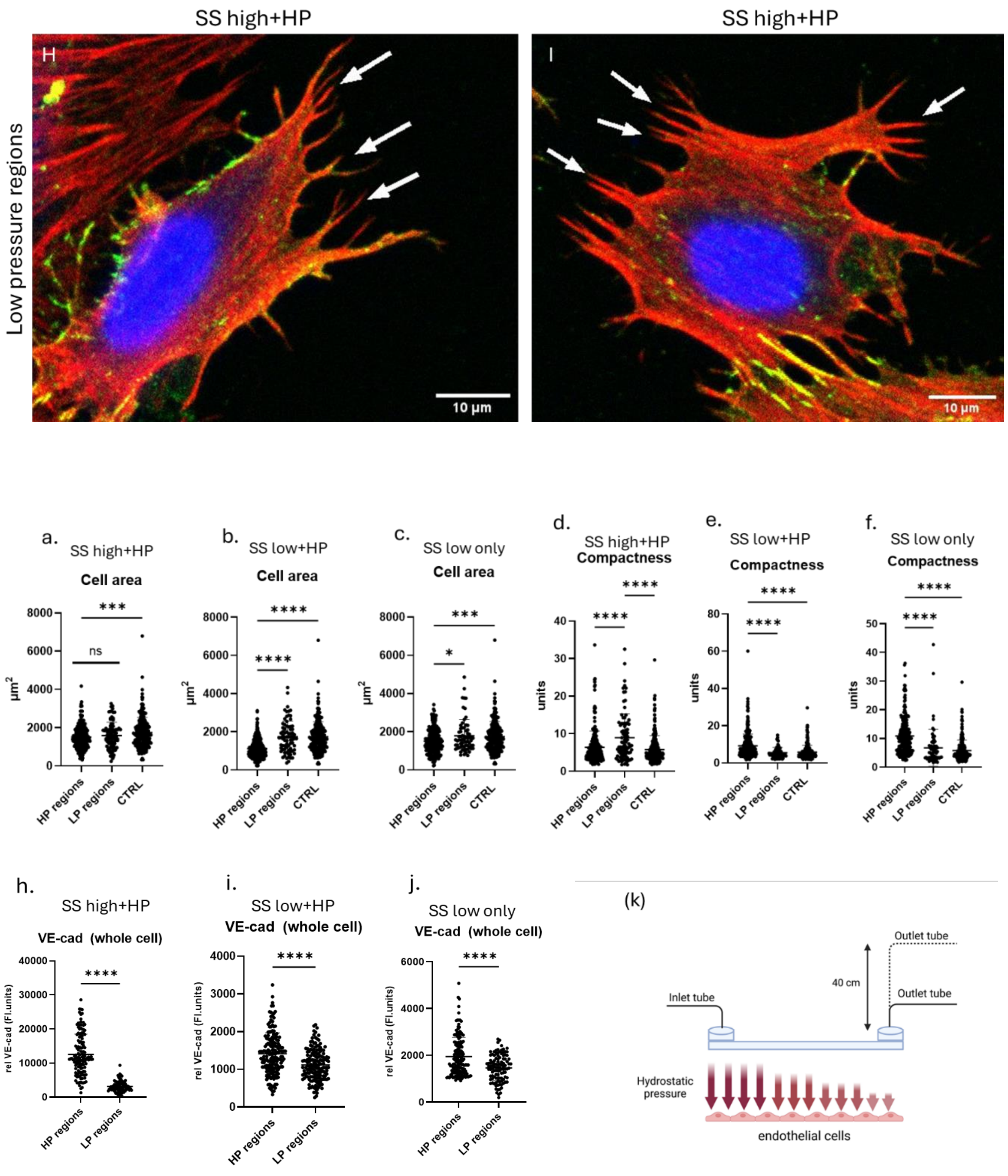


**Figure S3. The varying impact of elevated hydrostatic pressure in the high-pressure and low-pressure regions of the microfluidic channel.** Low-pressure regions (A-C) demonstrated less confluent cell density with increased gaps between cell clusters compared to the high-pressure regions (D-F) which showed reduced gaps between cells. (A,D) Cells exposed to SS high+HP demonstrated a significant reduction in VE-cadherin finger formation and VE-cadherin F.I. (h) in the low pressure regions compared to the high-pressure regions although there was no change in average cell area (a). Cell clusters in the low-pressure region also demonstrated tricellular VE-cadherin junctions (yellow arrows in (A)) which was not observed in the high-pressure regions. In the low-pressure regions of (B,E) SS low+HP and (C,F) SS low only conditions, cells demonstrated a significant increase in the average cell area possibly due to increased cell spreading to form contacts with neighbouring cells. Cells in this region also lacked continuous VE-cadherin expression and increased VE-cadherin finger formation at junctions as evidenced by the reduction in relative VE-cadherin fluorescence intensity (I,j). (H,I) Cells in the low-pressure regions of SS high+HP showed multi-filopodial protrusions branching from thick actin fibres. (a-c) average cell area in the high-pressure (HP) and low-pressure (LP) regions, (d-f) compactness of cell in the HP and LP regions, (h-j) relative VE-cadherin fluorescence intensity in the HP and LP regions. (k) Schematic representation of the variable pressure experienced by the endothelial cells within the channel. N = 3, at least 50 cells were analysed per repeat. Flow direction: Top to bottom. White arrows: multi-filopodial protrusions. White arrow heads: single filopodial protrusions. Yellow arrows: tricellular junctions. Mag = 25x

SS high+HP conditions did not cause any significant difference in cell area and eccentricity, but the cells were more irregularly shaped as evidenced by the increased compactness in Figure S3 (d). There was an increased number of multi-filopodial protrusions at cell edges (white arrows in Figure S3 (H,I)) in the low-pressure regions compared to the high-pressure region cells, which mostly exhibited single filapodial protrusions (white arrowheads in Figure S3D). Interestingly, in the low-pressure regions of SS high+HP, cell clusters demonstrated continuous junctions with many cells forming tri-cellular junctions (yellow arrows in Figure S3A). Such continuous junctions were not observed in the high-pressure regions where cells mostly formed VE-cadherin fingers at junctions. In contrast to the SS high+HP, for the SS low+HP and SS low only conditions, cells in the low-pressure regions demonstrated a significant increase in cell area, possibly due to the low cell density in this region, prompting cells to spread more to form cell-cell contacts. In both these conditions, there was also an increase in the number of filopodial protrusions formed (white arrows in Figure S3B and S3C) in the low-pressure regions similar to that of cells in the low-pressure regions of SS high+HP and there was a decrease in the average VE-cadherin F.I. in all three conditions as shown in Figure S3 (h,i,j). These results demonstrate the significant impact of local pressure drop across the microfluidic channel on VE-cadherin and actin dynamics. Cell density also influenced the patterning of VE-cadherin fingers in the SS high+HP conditions with cells in the high pressure regions exhibiting more prominent VE-cadherin finger structures compared to the low pressure regions as shown in Figure S4 below.

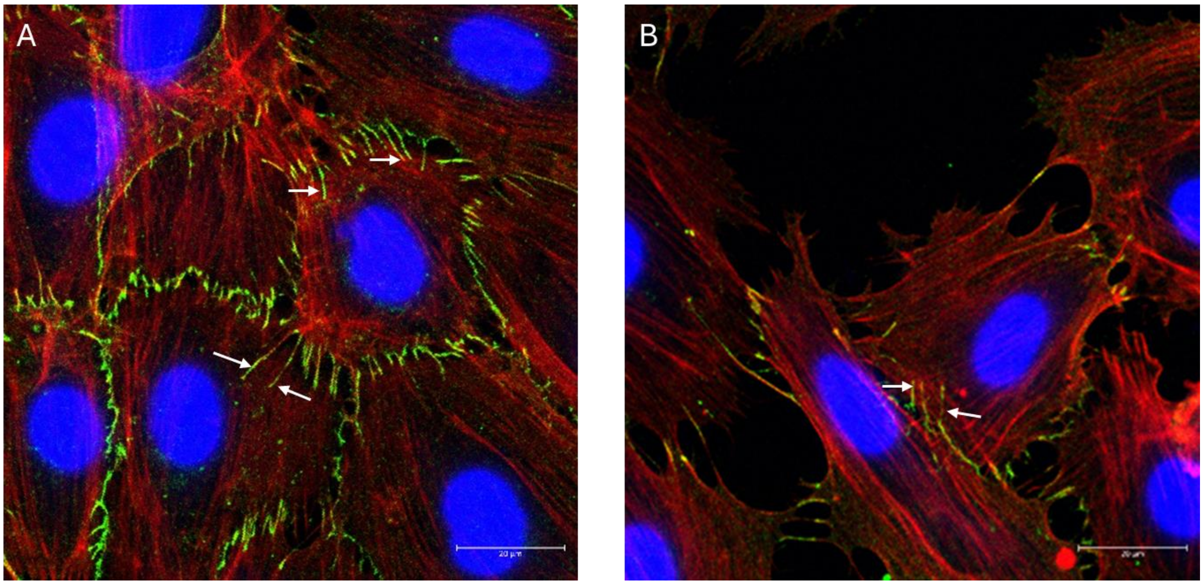

**Figure S4. Cell density dependent differential VE-cadherin finger formation.** (A) Areas of high cell density in SS high+HP conditions demonstrated thick, mature VE-cadherin fingers (white arrows) that extending well into the neighbouring cells cytoplasm. (B) In regions of low cell density, very few VE-cadherin fingers were observed even in the cells that expressed multiple, long filopodia protrusions that made contacts with the neighbouring cells. N = 3, at least 50 cells were analysed per repeat. Flow direction: Top to bottom. White arrows- VE-cadherin fingers. Mag = 63x

### 1.4 Peizo-1 inhibits lamellipodia and filopodial formation at elevated hydrostatic pressure conditions

As shown in Figure S6 (A) and (C), the extensive lamellipodial (white arrow heads) and filopodial protrusions (white arrows) observed in SS high+HP and SS low+HP conditions were inhibited upon blocking piezo-1 and instead exhibited a predominantly a thick, radial, actin meshwork around the cell membrane (green arrows in Figure S6 (B,D)).

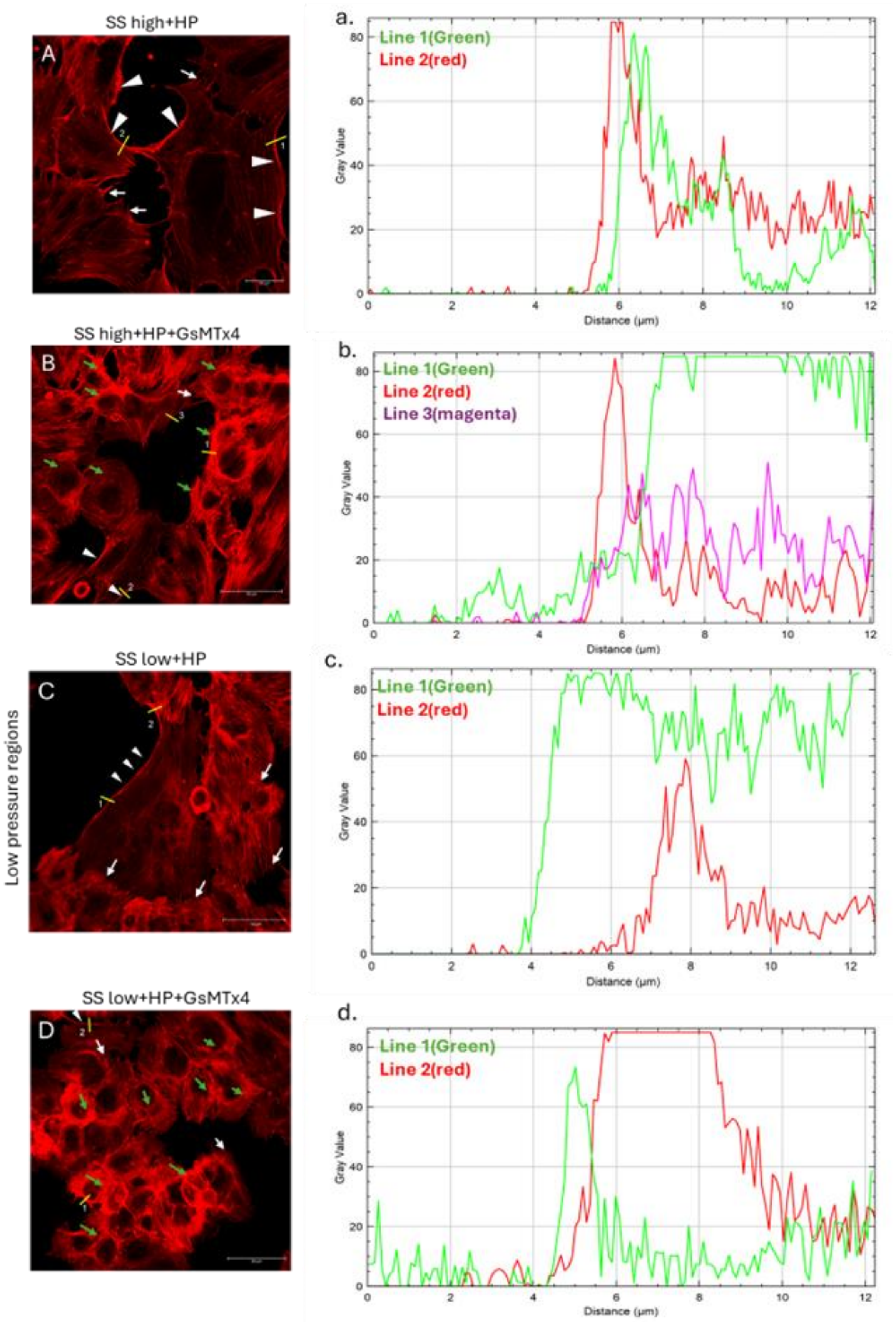


**Figure S5. Influence of piezo-1 inhibitor on the influence of elevated hydrostatic pressure on actin remodelling.** (A) SS high+HP demonstrated several regions with distinct lamellipodia formation (white arrow heads) as well as thick filopodial protrusions (white arrows). Line plots (a, b) demonstrate that the lamellipodial structures are surrounded either by thick fibres (line 1: green plot) or a mesh of thin fibres that end in filopodia structures (line 2: red plot). (B) In the presence of piezo-1 inhibitor GsMTx4, cells predominantly exhibited thick, randomly aligned radial, actin fibres as indicated by the green arrow heads with few cells exhibiting thin filopodial and lamellipodial protrusions. Line plots indicate the high-density mesh of cortical actin fibres in the polygonal shaped cells (line 1: green plot). The cells that demonstrated lamellipodia structures were surrounded by low density fibres (line 2: red plot) while some cells demonstrated thick array of actin fibres that did not form lamellipodial structures (line 3: magenta plot). (C) SS low+HP cells contained thick, aligned stress fibres ending in protrusions (white arrows) and lamellipodia (white arrow heads) with very few cells exhibiting cortical actin. Line 1(green plot) shows lamellipodia formation in a dividing cell. The plot profile shows decreased density of actin fibres adjacent to the lamellipodia structure. Line 2 (red plot) shows that the same dividing cell can have lamellipodial structures that are supported by dense network of aligned stress fibres. In the SS low+HP conditions, only cells in the low-pressure region which had low cell numbers exhibited thick filopodial and lamellipodial protrusions while the high-pressure region cells mostly exhibited radial actin fibre arrangements. (D) Similar to that of the SS high+HP, the addition of GsMTx4 to the cells in the SS low+HP caused majority of cells to predominantly develop thick, randomly aligned, cortical actin fibres (green arrows) with very few cells exhibiting lamellipodia (white arrow heads) and filopodia (white arrows). Line 1 (green plot) represents a thick cortical actin mesh while Line 2 (red plot) represents a thick lamellipodia formation. Flow direction: Top to bottom. Mag = 40x

### 1.5 VE-cadherin and actin co-localisation in 24h exposure to elevated hydrostatic pressure

In the SS high+HP conditions, there was increased co-localisation of VE-cadherin and actin especially at the serrated VE-cadherin junctions at the front or rear of the cells (Figure S6(A), line plot 1) whereas at the sides, cells demonstrated continuous stable VE-cadherin junctions that co-localised with actin (Figure S6(A), line plot 2) and a few cells demonstrated little to no VE-cadherin at the junctions (Figure S6(A), line plot 3). The results were in stark contrast for SS low+HP prolonged exposure conditions. There was little to no co-localisation of VE-cadherin/actin- neither at the side junctions of the cells (Figure S6(B), line plot 1) nor at the serrated junctions at the front or back of the cells (Figure S6(B), line plot 2). Many cells demonstrated only dotted VE-cadherin junctions (Figure S6(B), line plot 3) as most of the VE-cadherin were concentrated in the cytoplasm as opposed to the membrane.

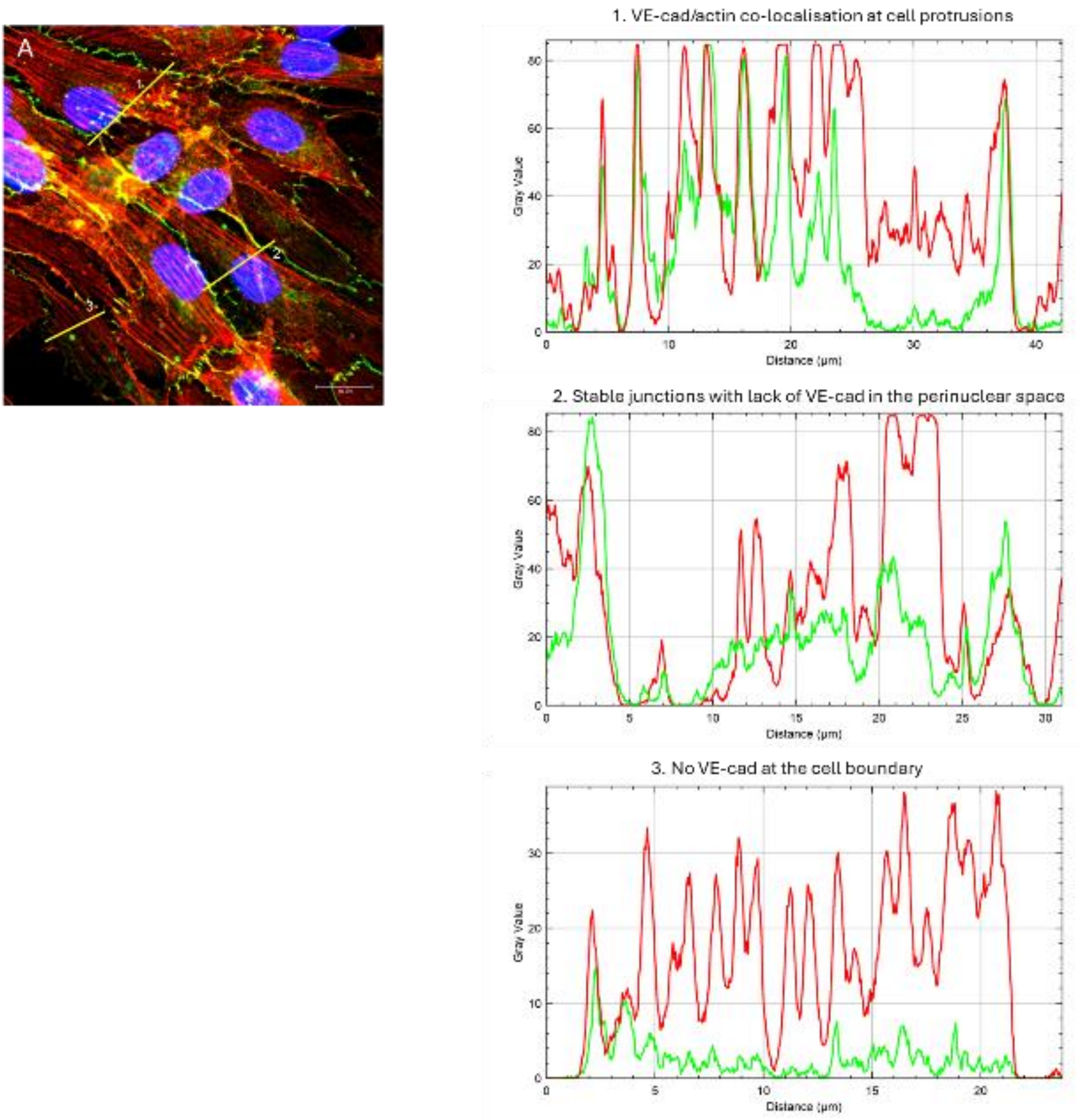

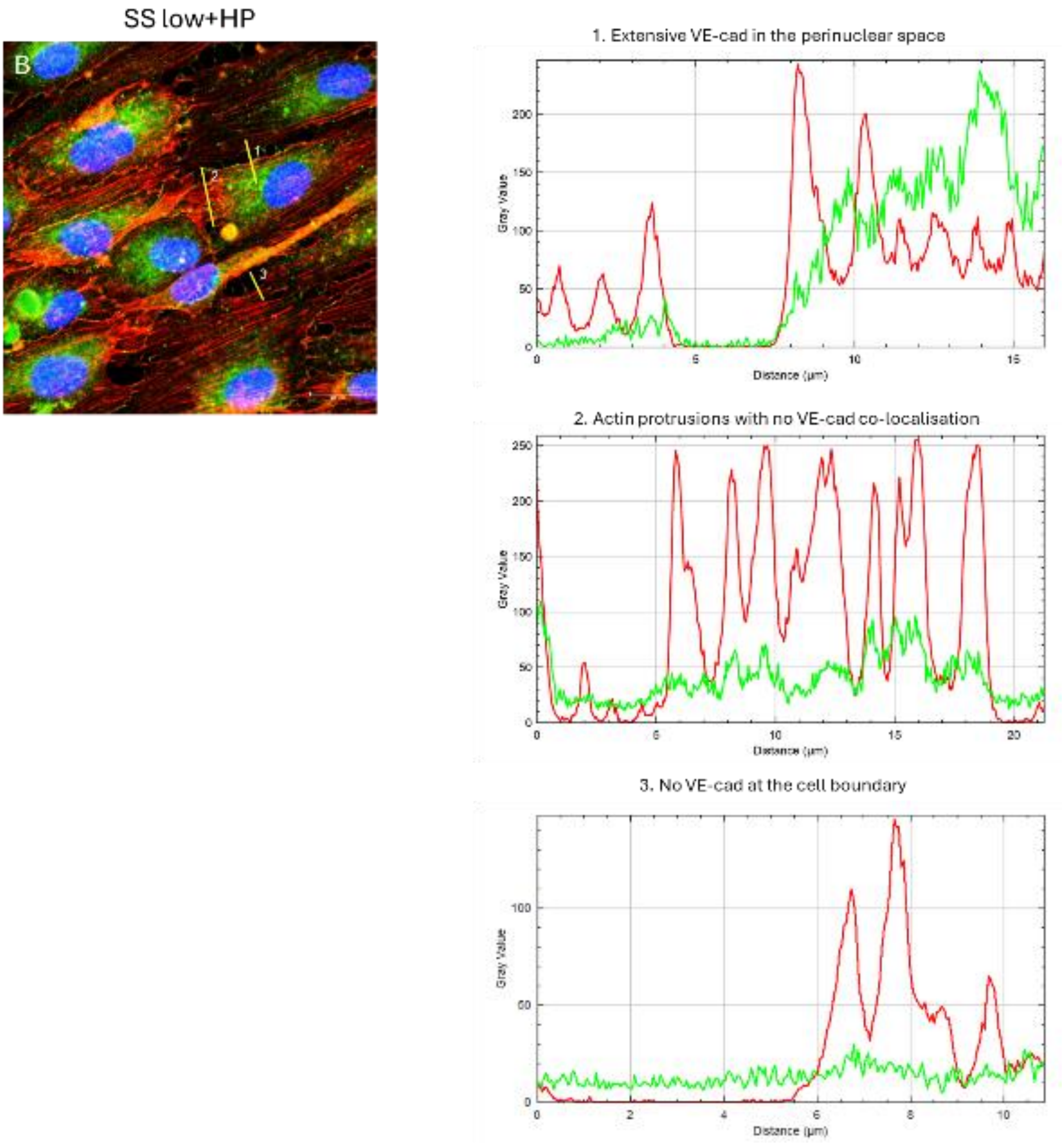


**Figure S6. Long-term exposure to elevated hydrostatic pressure leads to decreased VE-cadherin finger formation in SS low+HP conditions.** (A) SS high+HP demonstrated VE-cadherin finger formation at cell-cell junctions at the front and rear of cells (line plot 1) as well as continuous VE-cadherin junctions (line plot 2) at the sides of the cell (B) SS low+HP conditions demonstrated extensive VE-cadherin localisation in the cytoplasmic perinuclear space (line plot 1) and there was a lack of VE-cadherin co-localisation at the actin protrusions at the membrane (line plot 2). Note that in both conditions, there were many cells which lacked continuous VE-cadherin at the cell-cell junctions. Red plot line: actin, Green plot line: VE-cadherin. Flow direction: Left to right. Mag = 63x
